# Peripheral sgp130-mediated *trans*-signaling blockade induces obesity and insulin resistance in mice via PPARα suppression

**DOI:** 10.1101/2020.09.24.309716

**Authors:** Tali Lanton, Orr Levkovitch-Siany, Shiran Udi, Joseph Tam, Rinat Abramovitch, Sharon Perles, Evan Williams, Jacob Rachmilewitz, Uria Mor, Eran Elinav, Dirk Schmidt-Arras, Ateequr Rehman, Philip Rosenstiel, Anastasios Giannou, Samuel Huber, Stefan Rose-John, Eithan Galun, Jonathan H. Axelrod

## Abstract

IL-6 signaling via its receptor (IL-6R) and co-receptor (gp130) performs multiple roles in regulating metabolic homeostasis. However, gp130 is also expressed systemically in a soluble form (sgp130), which limits soluble IL-6 receptor (sIL-6R)-mediated signaling – also called *trans*-signaling. Here we find that transgenic peripheral sgp130-mediated *trans*-signaling blockade induces mature-onset obesity, while differentially affecting age-dependent behavioral determinants of energy expenditure. In youth, *trans*-signaling blockade increases feeding associated with reduced leptin sensitivity but increases energy expenditure to maintain metabolic balance. In aging, reduced physical activity predisposes mice to adiposity, adipose tissue macrophage recruitment, hepatosteatosis, hyperglycemia, and insulin resistance. Mechanistically, *trans*-signaling blockade reduces hepatic Stat3 phosphorylation and suppresses PPARα, associated with miR-21 upregulation, while pharmacological activation of PPARα prevents obesity and hepatosteatosis, and rescues insulin sensitivity. Together these experiments reveal a role for peripheral IL-6 *trans*-signaling in metabolic homeostasis and provide clinical significance to elevated sgp130 levels found in some obese and diabetic patients.

## INTRODUCTION

Substantial evidence indicates that IL-6 is crucial for maintaining metabolic homeostasis affecting multiple peripheral organs. Thus, mice with total IL-6 deficiency develop mature-onset obesity, characterized by increased adiposity and hepatic steatosis accompanied by glucose intolerance, hyperinsulinemia, and systemic insulin resistance (Matthews et al., 2010; Wallenius et al., 2002). Similarly, selective disruption in mice of IL-6 signaling in the liver by targeted ablation of the IL-6 receptor (IL-6R) in hepatocytes increases inflammation and reduces insulin sensitivity (Wunderlich et al., 2010). Physical exercise has been found to induce elevation of muscle-derived IL-6 up to 100-fold in the circulation (Pedersen & Febbraio, 2008), and is associated in humans with the enhancement of insulin-stimulated glucose disposal and increased insulin sensitivity in the peripheral tissues (Carey et al., 2006). Moreover, long-term blockade of IL-6 signaling by ACTEMRA® (tocilizumab), a recombinant humanized anti-IL-6 receptor (IL-6R) antibody, is associated with significant increases in body weight and serum lipid and cholesterol levels and abolished the exercise-induced reduction in visceral adipose tissue mass (Kawashiri et al., 2011; Wedell-Neergaard et al., 2018).

Together with its systemic effects, IL-6 and its receptor are also constitutively expressed in the central nervous system (CNS) (Schobitz et al., 1993), where an increasing body of evidence points to its role in the regulation of metabolic homeostasis. IL-6-dependent pathways in the lateral parabrachial nucleus exerted control on body weight (Mishra et al., 2019) and in the hypothalamus, they contribute to reduce feeding and maintain peripheral glucose tolerance, especially in the face of leptin and insulin resistance in obesity (Timper et al., 2017). Thus, IL-6 interacts at both peripheral and central sites in the body, and these interactions, in general, appear to promote metabolic homeostasis (Jansson & Palsdottir, 2015).

Notably, the cognate IL-6 receptor is found in two forms, a membrane-bound form (IL-6R) and an abundantly expressed soluble form (sIL-6R), leading to multiple signaling configurations. In its *classical* signaling configuration, membrane-bound IL-6R, which is expressed on limited cell populations, combines with IL-6 to engage its ubiquitously expressed, transmembrane co-receptor, gp130, to initiate intracellular downstream signaling cascades, such as the JAK/STAT pathway (Heinrich et al., 2003). However, in its soluble-form, sIL-6R can form a complex with IL-6 and, in a mechanism called *trans-*signaling, initiate signaling in gp130 expressing cells, including in cells that do not express membrane-bound IL-6R (Rose-John et al., 2006). Importantly, gp130 is also produced in a truncated, soluble form (sgp130) that is normally expressed in the serum. There sgp130 functions to bind and neutralize IL-6/sIL-6R complexes, but not either protein alone, thus acting as a specific antagonist of *trans-*signaling (Jostock et al., 2001; Narazaki et al., 1993). It has been postulated that the abundant sgp130 and sIL-6R in the serum act as a natural buffer to localize and limit the potentially deleterious effects of excessive IL-6 *trans-*signaling, such as found in some inflammation-driven pathologies (Rose-John et al., 2006). In contrast, in the hypothalamus, *trans*-signaling underlies a crucial IL-6-mediated, leptin-independent mechanism that functions to suppress food-intake and maintain glucose tolerance especially in obesity (Timper et al., 2017).

However, the importance of *trans*-signaling in peripheral, metabolism-related IL-6 interactions remains unclear. In accordance with its role in immune activation, IL-6 *trans*-signaling is important for the obesity-associated recruitment of macrophages into adipose tissue in humans and also for macrophage chemotaxis *in vitro* (Kraakman et al., 2015), and as such would be expected to support glucose intolerance and insulin resistance in diabetes. Importantly, peripheral *trans*-signaling blockade can be imposed by high-level systemic expression of a recombinant sgp130 protein combined as a fusion protein with the fragment crystallizable (Fc) domain of IgG, called sgp130Fc (Jostock et al., 2001), which has also been engineered into a transgenic mouse model (herein referred to as *‘sgp130Fc mice’*) (Rabe et al., 2008). In this model, *trans*-signaling blockade in young, HFD-challenged mice prevents adipose tissue-associated macrophage (ATM) recruitment, but neither relieves nor exacerbates HFD-induced weight gain, hepatic steatosis, or insulin resistance in the young mice (Kraakman et al., 2015). Similarly, in young mice on a normal chow diet, ectopic sgp130 overexpression does not significantly alter body mass and does not affect glucose tolerance or energy expenditure in comparison to wild type controls (Kraakman et al., 2015). Paradoxically, however, elevated systemic sgp130 levels have been found in multiple studies to be associated with obesity, diabetes, and insulin resistance in adult patients (Nikolajuk et al., 2010; Weiss et al., 2013; Zuliani et al., 2010), although not in obese children or adolescents (De Filippo et al., 2015). Moreover, in patient populations with coronary artery disease (CAD) or at high risk for CAD, elevated serum levels of sgp130 were found in older individuals carrying the gp130 polymorphism, G148C, and were also associated with diabetes and increased body mass index (Wonnerth et al., 2014). This polymorphism has been shown to impair the functionality of gp130 (Benrick et al., 2008), thus suggesting that sgp130 polymorphisms may pose a potential genetic predisposition to co-morbidities of metabolic syndrome.

These observations suggest that peripheral *trans*-signaling may play an important role in the maintenance of metabolic homeostasis, but that aging is a critical factor in its presentation. In order to test this hypothesis and to unfold its potential mechanism(s), we raised sgp130Fc mice to the age of 14 months and assessed body composition, weight gain, blood glucose, and metabolic and behavioral determinants of energy expenditure in young *versus* aged mice. Our observations reveal that, in aging, blockade of peripheral *trans*-signaling induces striking behavioral and metabolic aberrations, including mature-onset obesity, glucose intolerance, and insulin resistance. We further identify the suppression of the Stat3-regulated miR-21/PPARα axis as a mechanism underlying some aspects of this phenotype.

## RESULTS

### Inhibition of *trans*-signaling induces mature-onset obesity on a normal chow diet

Between the ages of about 5 and 6 months, the body weights of male sgp130Fc mice begin to diverge from their WT littermates, becoming visibly and significantly overweight after 7 months of age (Figures 1A,B). The increase in body fat in mature sgp130Fc mice was shown by MRI imaging analysis (Figures 1C,D), by gonadal fat pad dissection upon sacrifice at 14 months (Figure 1E), and by echoMRI analysis (Figures 1F and S1B). The increase in body weights was due largely to fat accumulation, which, according to MRI analysis, increased by about 40 percent in sgp130Fc mice at 14 months of age compared to their WT littermates, with roughly equal distribution between subcutaneous and abdominal fat mass (Figures 1D,E). Accordingly, prior to major bodyweight divergence the levels of the adipose-secreted hormone leptin were unchanged in young sgp130Fc mice aged 2-7 months relative to controls, in agreement with previous observations (Kraakman et al., 2015), but increased significantly in comparison to their WT littermates by 12 months of age commensurate with their increased fat mass (Figure 2A).

**Figure 1.**
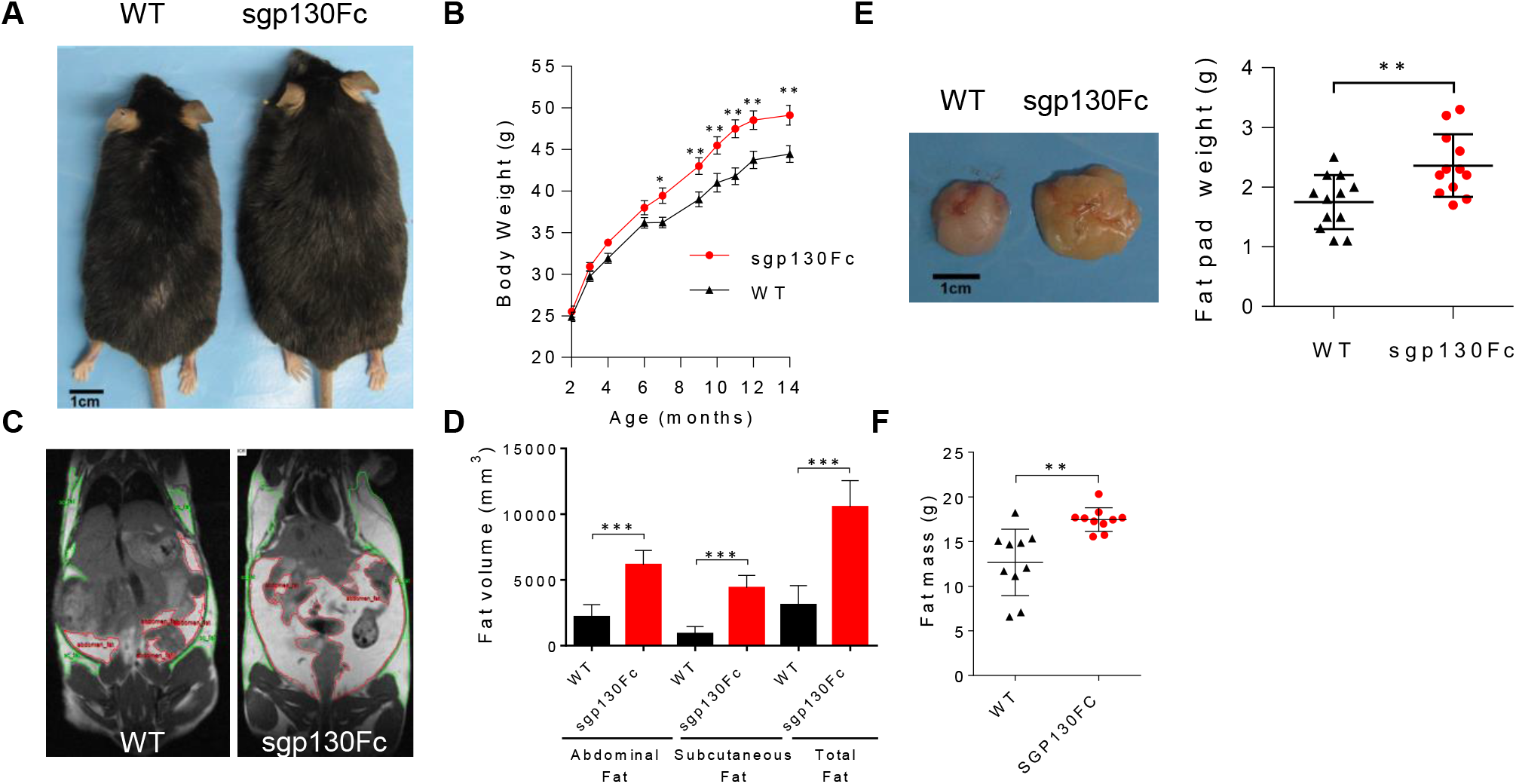
Sgp130-mediated *trans*-signaling blockade induces mature-onset obesity in mice on a normal chow diet. (A) Photographs showing representative male, sgp130Fc and WT mice at 14 months of age. (B) Body weight progression from 2 to 14 months of age in male, sgp130Fc and WT mice (n=9-20). The primary observation repeated in 4 independent experiments in male littermates and once in both male and female non-littermate mice. (C) Representative coronal MRI images of sgp130Fc and WT mice at 10 months. (D) Quantification of subcutaneous and abdominal fat tissue in mice from MRI images (n=4-5) (E) Photographic image of representative gonadal fat pads and quantification of fat pad mass (right) at 14 months (n=12). (F) EchoMRI quantification of fat mass at 14 months (n=9-10). Data are mean ± SD (E-F), or mean ± SEM (B). \*\**P* < 0.01, and\*\*\**P* < 0.001, by two tailed, Student’s t test. (See also Figure S1.)

**Figure 2.**
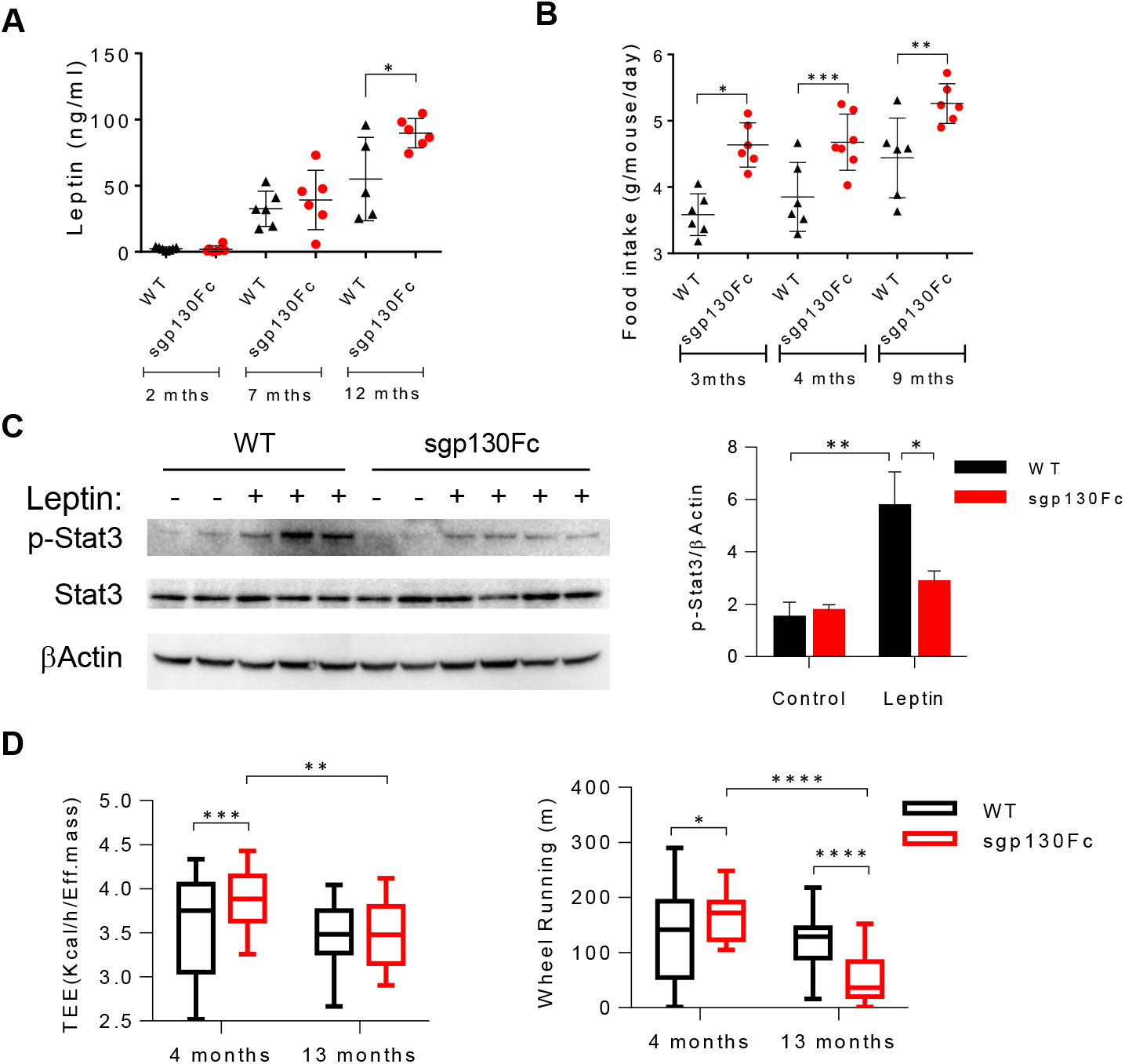
Peripheral *trans*-signaling blockade induces metabolic and behavioral changes in mice. (A) Serum leptin concentrations at 2, 7, and 12 months (mths) (n=5-6). (B) Daily food-intake at 3, 4 and 14 months of age (n=6-7). Data represent mean food intake per mouse/cage per day averaged over a 2-week period. (C) Western blot analysis and quantification (right) of Stat3 phosphorylation in the hypothalamus of 3.5 month old mice treated with leptin or carrier control (n=2-4). (D) Metabolic cage assessment of dark cycle total energy expenditure (TEE) and voluntary physical activity (Wheel Running) of mice at 4 and 13 months of age (n=12). Data are represented as mean ± SD. \**P* < 0.05, \*\**P* < 0.01, \*\*\**P* < 0.001, and\*\*\*\**P* < 0.0001 by two-tailed, Student’s t test (G,H), or by two-way ANOVA (C,D). Significant interactions were reported for leptin induced p-STAT3 and genotype (*P*=0.0193), and for genotype and age for TEE (**P=0.0024) and Wheel Running (\*\*\*\**P* < 0.0001). (See also Figure S1.)

### Etiology of weight gain in sgp130Fc mice

To elucidate the underlying causes of body weight gain in sgp130Fc mice we measured daily food intake and assessed energy expenditure using metabolic cages. Beginning from a young age (3 months) and prior to the onset of obesity, food intake by sgp130Fc mice was consistently increased by about 30 percent compared to WT littermates and remained high at older ages (Figure 2B). Interestingly, the elevated food intake levels, suggestive of leptin resistance, were consistent with the hyperleptinemia in older sgp130Fc mice, but were less clear for younger mice, which did not display increased serum leptin levels (Figure 2A). However, Western blot analysis following leptin injection revealed a strong reduction in leptin-induced Stat3 phosphorylation levels in the hypothalamus of young sgp130Fc mice (Figure 2C), thus indicating that prior to weight gain, leptin-sensitivity is reduced in younger sgp130Fc mice as well. This suggests that elevated peripheral *trans*-signaling blockade interferes with *central* leptin signaling, perhaps accounting in part for the increased food intake in the young sgp130Fc mice, and later on in weight gain in the aged animals.

Metabolic cage analyses revealed that young (4 months) sgp130Fc mice also display overall increased physical activity, particularly voluntary nocturnal activity including wheel running, and elevated energy expenditure relative to WT littermates (Figures 2D and S1C). However, while nocturnal energy expenditure and voluntary physical activity decreased only marginally over time in WT controls, in sgp130Fc mice both parameters decreased substantially such that by 13 months of age energy expenditure and wheel running were reduced, respectively, by 11 and 67 percent relative to their younger counterparts (Figures 2D, and S1C,D). Accordingly, sgp130Fc mice also displayed lower nocturnal respiratory quotient (RQ) that decreased further with age and elevated utilization of fat versus carbohydrate oxidation that decreased with age (Figure S1D). Statistical analyses confirmed the strong interaction between the metabolic phenotype of the sgp130Fc mice and age (*P*<0.001), indicating that age is likely a substantial factor in the mechanism of weight gain in the sgp130Fc mice (Figure 2D and S1D).

Importantly, the increased adiposity in sgp130Fc mice appeared to be due to adipocyte hypertrophy since, as assessed by H&E staining of white adipose tissue (WAT) biopsies, a significant increase in the mean adipocyte cross-sectional area emerged concurrently with the onset of divergence in body weights in the mature mice (Figures 3A,B and S2A). In addition, ATM recruitment and crown-like structures characteristic of adipose tissue in obesity (Makki et al., 2013) increased in aged sgp130Fc mice compared to controls, as quantified by anti-F4/80 immunostaining (Figures 3C,D) and confirmed by FACS analysis (Figure 3E). The increase in ATM in sgp130Fc mice also appears to be age-dependent since the opposite was reported for younger sgp130Fc mice upon challenge with an HFD (Kraakman et al., 2015).

**Figure 3.**
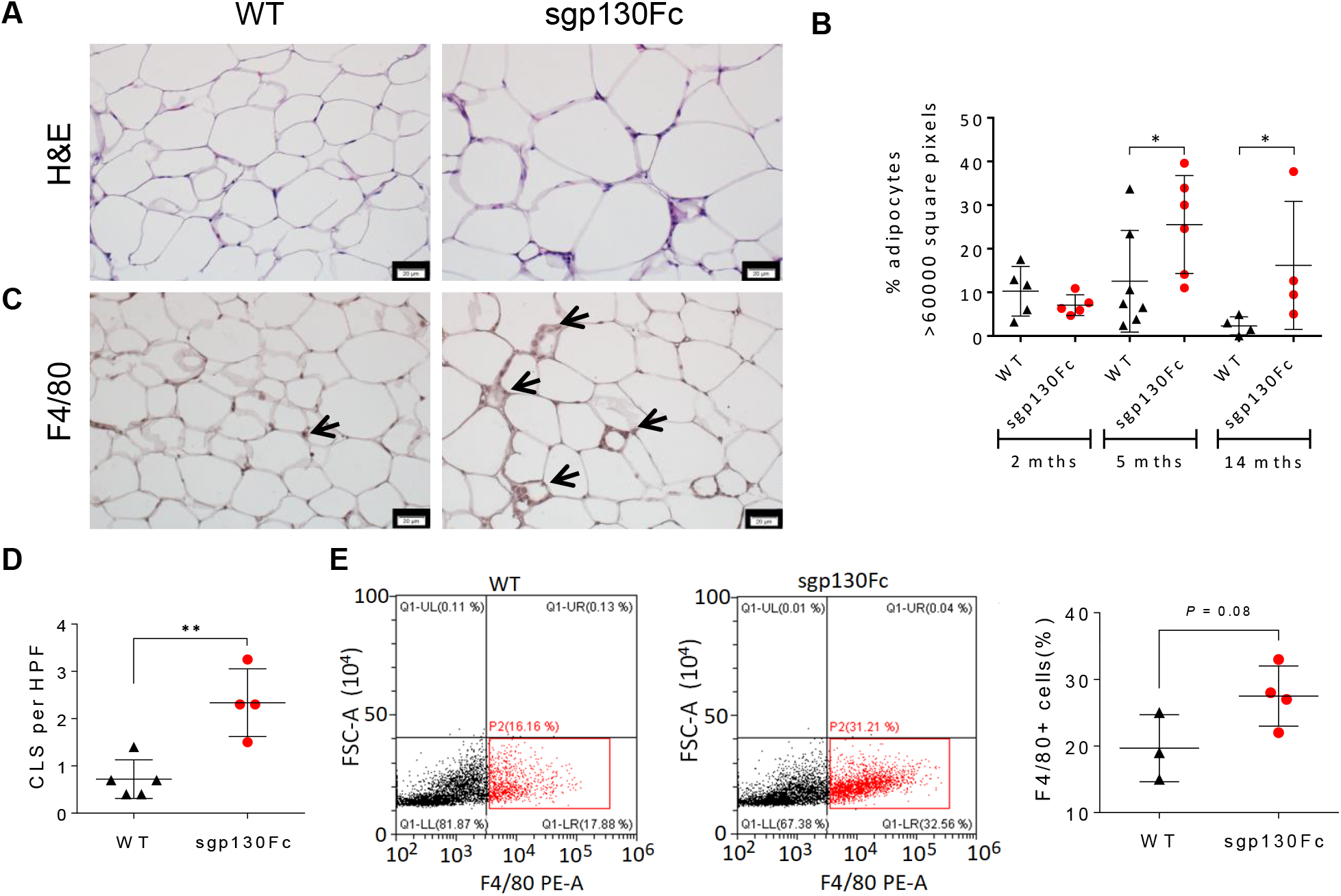
IL-6 trans-signaling blockade induces adipocyte hypertrophy and ATM recruitment. (A) H&E stained adipose tissue sections in sgp130Fc mice and WT littermates at 14 months of age. Scale bars, 20μm. (B) Quantification of adipocytes with cell cross-sectional area >6000 square pixels by ImageJ, (n=4-7). (C) Photomicrograph of F4/80 immunostaining of adipose tissue thin sections showing adipose tissue-associated macrophage (ATM) (red) in crown-like structures (CLS) (arrows) at 14 months. Scale bars, 20μm. (D) Quantification of F4/80+ CLS per high power field (HPF) in adipose tissue thin sections (c), (n=4). (E) Representative flow cytometry plots (left) and quantification (right) of F4/80^+^ cells in adipose tissue of 11-month old mice, (n=3-4). F4/80+ cells were quantified as per the bottom right quadrant (gate P2) (red). Data are represented as mean ± SD. * P < 0.05, and ** P < 0.01 by two-tailed, Mann Whitney test (B), or two-tailed, Student’s t test (D, E). (See also Figure S2.)

### Sgp130Fc mice display hyperglycemia, peripheral insulin resistance and hepatosteatosis in the absence of hepatic inflammation

Because obesity and hyperglycemia are hallmarks of the metabolic syndrome, we next assessed the mice for glucose tolerance and insulin resistance at both early and late phases of weight gain. Sgp130Fc mice displayed fasting hyperglycemia (Figure 4A) and significant glucose intolerance that was exacerbated with age (Figure 4B and S3A). The overweight sgp130Fc mice also displayed insulin tolerance test curves distinctive for peripheral insulin resistance (Figure 4C), and both fasting and post-prandial hyperinsulinemia compared to WT littermates (Figure 4D). Western blot analysis of ribosomal protein S6 phosphorylation in peripheral tissues (liver, muscle, and adipose) in response to injected insulin in 10-month old sgp130Fc mice confirmed the presence of peripheral insulin resistance (Figure S3B-D). Thus, hyperglycemia in sgp130Fc mice is likely an outcome of peripheral insulin resistance consequent to the adiposity observed in the mature sgp130Fc mice.

**Figure 4.**
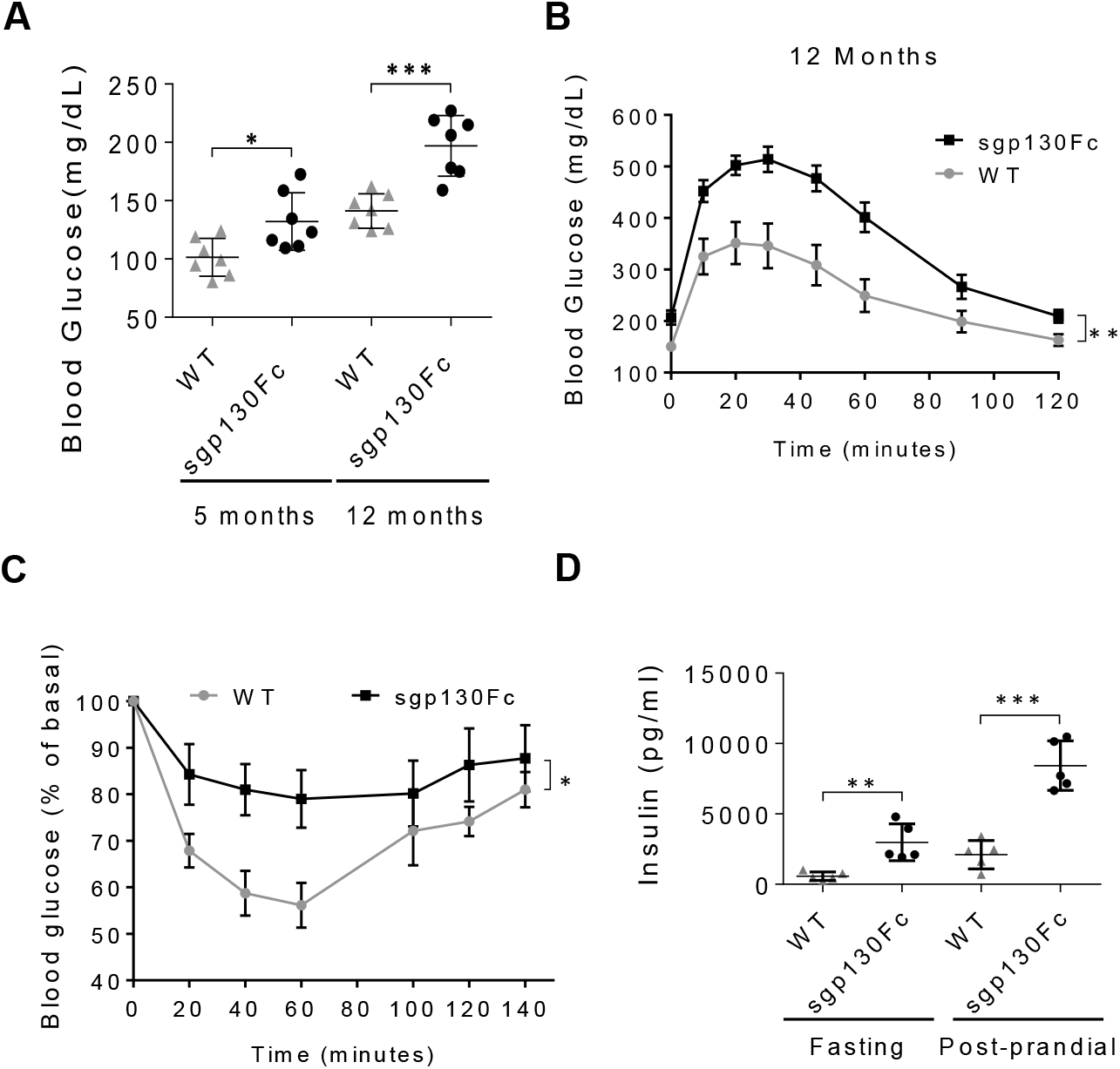
Inhibition of IL-6 *trans*-signaling induces glucose intolerance and peripheral insulin resistance. (A) Fasting glucose levels in sgp130Fc and WT littermates (n=7). (B) Glucose levels before and after glucose challenge in sgp130Fc versus WT mice at 5 (left) and 12 (right) months of age (n=7). (C) Blood glucose levels in fasting 12-month-old sgp130Fc and WT mice at indicated times following insulin injection (i.p.) (n=7). (D) Fasting and postprandial serum insulin levels in sgp130Fc and WT littermates at 13 months of age (n=5). Data are represented as mean ± SD (A,D), or mean ± SEM (B,C). * *P* < 0.05, ** *P* <0.01, and *** *P* <0.001 by two-tailed Student’s *t* test. (B,C) * *P* < 0.05, ** *P* <0.01 for comparisons of area under the curve by two-tailed Student’s *t* test. (See also Figure S3.)

Since peripheral insulin resistance is also strongly associated with NAFLD and NASH (Gastaldelli, 2017), we determined whether long-term ectopic sgp130Fc overexpression affected liver morphology and function by assessing lipid accumulation and inflammation in mature mice. Analysis of lipid accumulation by oil red O (ORO) staining showed that sgp130Fc livers at 14 months of age harbored frank steatosis with strikingly larger fat vacuoles compared to WT littermates, with initial signs of steatosis appearing in the mice already at 5 months of age (Figures 5A and S2B). These observations appear to indicate that elevated sgp130 recapitulates many phenotypic aspects previously reported for IL-6 *knockout* (IL-6^-/-^) mice, including obesity, hyperglycemia and hepatic steatosis (Matthews et al., 2010; Wallenius et al., 2002). Curiously, however, in our hands, while IL-6^-/-^ mice displayed significant hepatic steatosis in comparison to littermates, they did so without obvious weight gain or hyperglycemia (Figure S4), suggesting that the effect of elevated sgp130 may be more complex than total IL-6 ablation.

**Figure 5.**
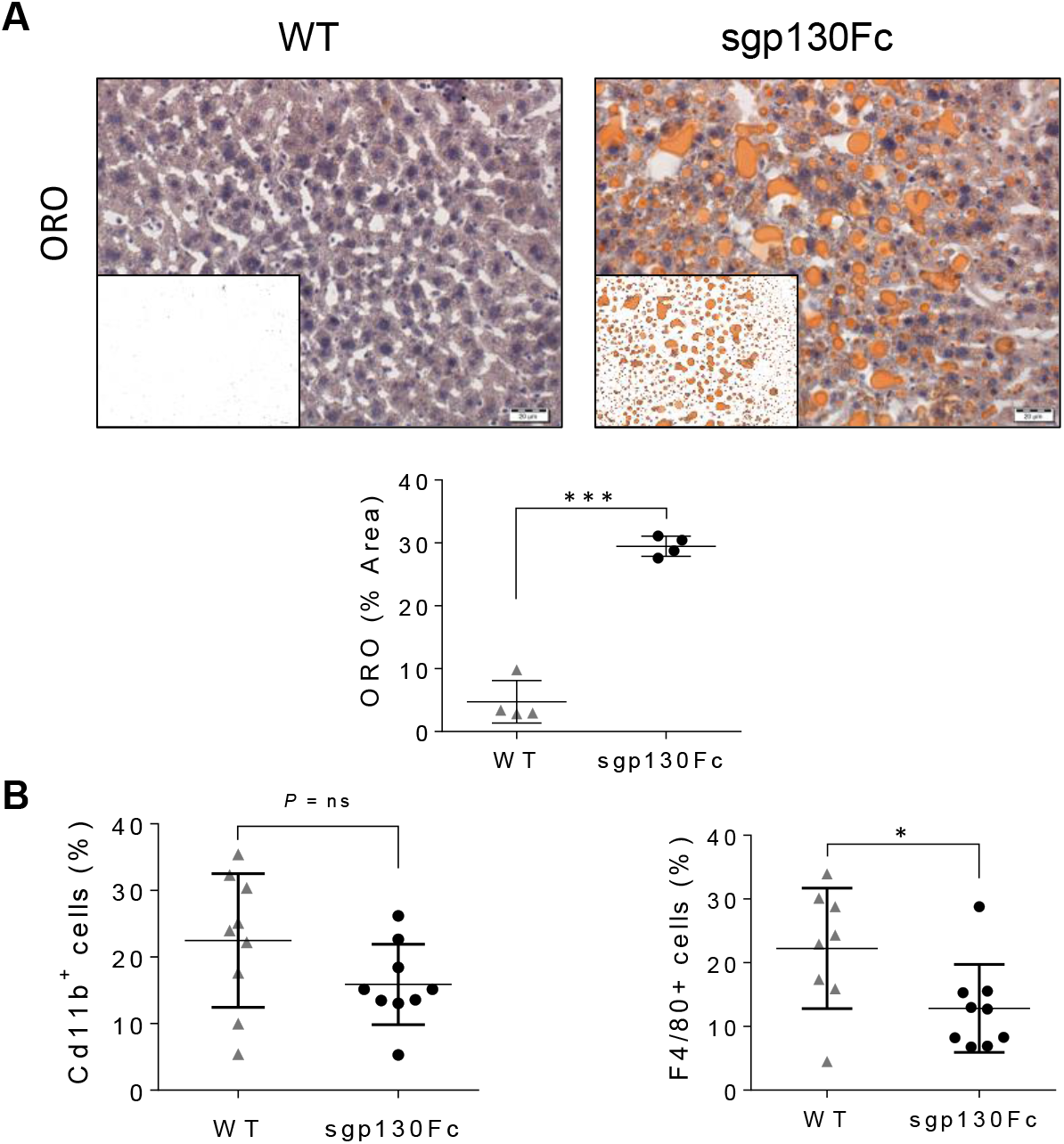
Inhibition of *trans*-signaling induces steatosis without hepatic inflammation. (A) Representative images showing fat accumulation by oil red O (ORO) staining in livers in sgp130Fc and WT littermates at the age of 14 months and ImageJ processed image (Inset). Scale bars represent 20 μm. (right) Quantification of ORO-stained area by Image J, (n=4). (B) Representative plots from flow cytometry of double-positive Cd45+/Cd11b+ and Cd45+/F4/80+ cells (upper right quadrant) in livers of sgp130Fc and WT littermates at 11 months of age and quantification (right), (n=8-9). Data are represented as mean ± SD. *P < 0.05, ***P<0.001 by two-tailed, Student’s *t* test. (See also Figures S2 and S5.)

A critical role of inflammatory signaling, associated in part with hepatic steatosis and increased Kupffer cells and mediated by tumor necrosis factor α (TNFα), has previously been identified in the induction of insulin resistance and glucose intolerance in mice with total IL-6 deficiency and also in mice with hepatocyte-targeted IL-6 receptor ablation (Matthews et al., 2010; Wunderlich et al., 2010). However, analysis of livers from aged (11-14 months old) sgp130Fc mice by flow cytometry, immunostaining, and qPCR revealed either reductions or no changes in the levels of monocytes/macrophages and inflammatory cytokine expression (Figures 5B and S5A). Thus, levels of Cd11b^+^ monocytes in the liver assessed by flow cytometry were unchanged between sgp130Fc and WT mice (Figure 5B), while populations of Kupffer cells and macrophages (F4/80) were either reduced (Figure 5B) or, as quantified by immunostaining, were unchanged (Figure S5B). Likewise, we observed no significant changes in levels of hepatic mRNAs encoding *Cd11b, F4/80, Cd68*, and *Tlr4*, or in the expression of inflammatory cytokines, *Tnfα, Il-10* and *Il-6* (Figure S5C). These observations suggest that in aged sgp130Fc mice hepatic inflammation-independent pathways may contribute to the development of insulin resistance and glucose intolerance.

### Suppression of hepatic PPARα signaling and miR-21 upregulation are associated with IL-6 *trans*-signaling blockade

IL-6 has been found to increase expression of hepatic nuclear transcription factor peroxisome proliferator-activated receptor alpha (PPARα), which is a master regulator of lipid metabolism, and has been found to be decreased in humans and mice with fatty liver disease (Liss & Finck, 2017; Montagner et al., 2016). Interestingly, *Pparα* expression can also be suppressed by miR-21, which targets PPARα expression in fatty liver disease (Kida et al., 2011; Loyer et al., 2016). Since miR-21 can be suppressed by signal transducer and activator of transcription 3 (Stat3) (Zhang et al., 2011), a primary mediator of IL-6 signal transduction, we hypothesized that suppression of PPARα associated with the reduction of Stat3 and elevation of miR-21 may be involved in the metabolic phenotype of our sgp130Fc mice.

Consistent with this hypothesis, we found that levels of phosphorylated Stat3 in livers of sgp130Fc mice were significantly reduced compared to WT littermates (Figure 6A). Additionally, we found that expression of miR-21 was consistently enhanced in livers of sgp130Fc mice (Figure 6B), while levels of miR-122, a liver-specific micro-RNA previously implicated in fat metabolism (Chai et al., 2017; Esau et al., 2006), did not change in a consistent manner between sgp130Fc and WT mice (Figure 6B). Analysis of *Pparα* mRNA and protein levels in livers of sgp130Fc and WT littermates at 2, 5, or 14 months of age showed significant reductions of *Pparα* mRNA expression in sgp130Fc mice relative to controls at 2 and 14 months (Figure 6C) and of PPARα protein as well, as shown at 5 months of age (Figure 6D). Importantly, compared to WT controls, *trans*-signaling blockade reduced mRNA levels of *Pparα* regulated genes, including *Cpt1, Apoe*, and *Fasn* (Hong et al., 2004; Vida et al., 2013) (Figure 6C).

**Figure 6.**
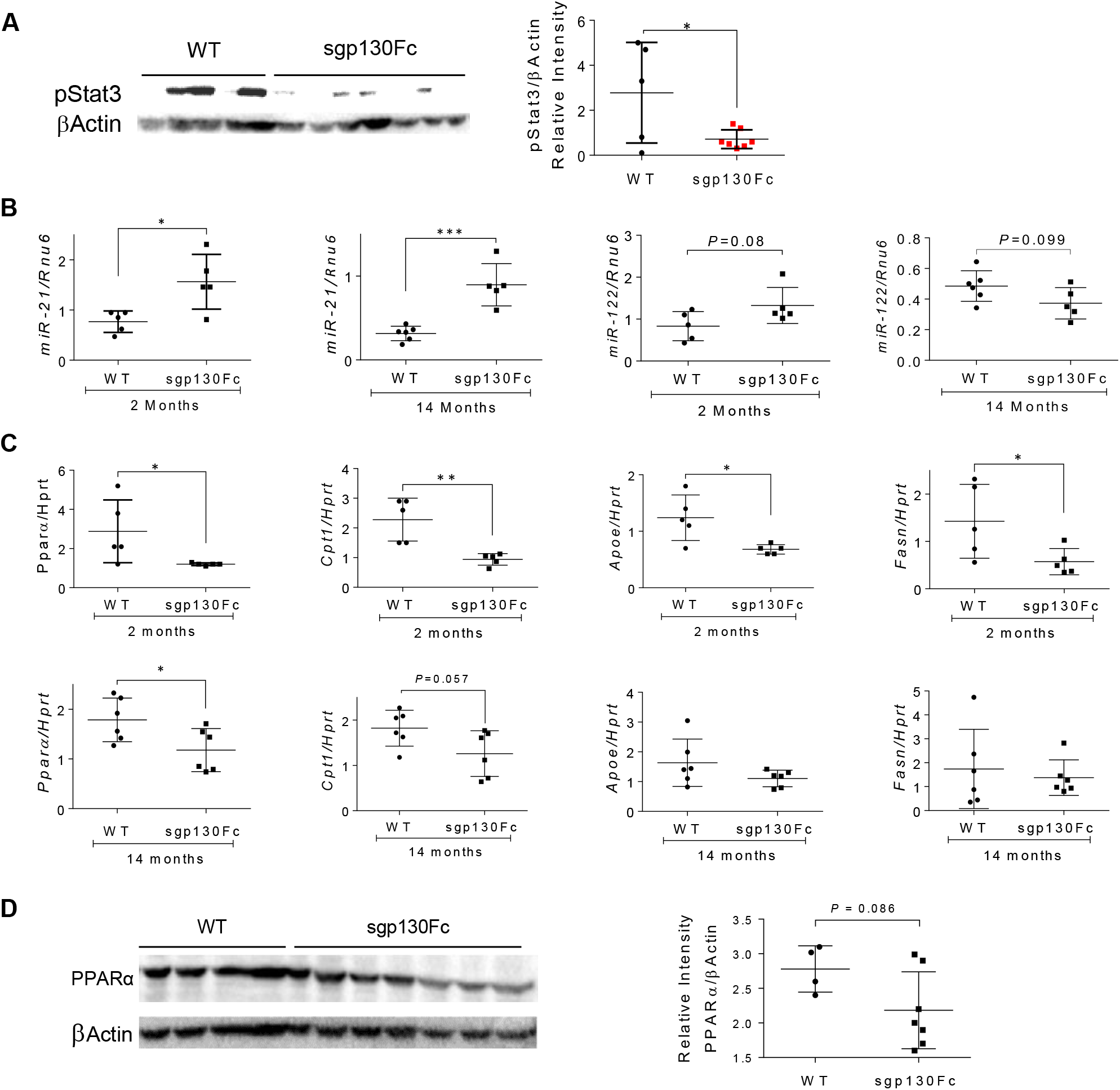
Suppression of hepatic PPARα and miR-21 upregulation are associated with IL-6 *trans*-signaling blockade. (A) Phosphorylated Stat3 and β-Actin protein levels by Western Blot analysis of liver samples from 5-month old sgp130Fc and WT littermates and quantification of band intensities (right). (n=5-7). (B) Hepatic miR-21 and miR-122 levels in sgp130Fc and WT littermates at 2 and 14 months of age by real time qPCR analysis, (n=5-6). (C) Real time qPCR analysis of hepatic mRNAs encoding PPARα, and PPARα targets: *Cpt-1, Apoe*, and *Fasn*, in sgp130Fc and WT littermates at 2 and 14 months of age (n=5). (D) Western blot analysis of PPARα and β-Actin protein levels in liver samples from 5 months old sgp130Fc mice and WT littermates with quantification of band intensity by Image J (right), (n=4-7). Data are represented as mean ± SD. * *P* < 0.05, ** *P* < 0.01, and *** *P* < 0.001 by two-tailed, Student’s *t* test. (See also Figure S6.)

To confirm the role of phosphorylated Stat3 in the regulation of PPARα, we examined the effect of hepatocyte-targeted Stat3 ablation in Alb-CreSTAT3^floxP^ (Stat3^Δhep^) mice (Inoue et al., 2004) on hepatic miR-21 and PPARα expression levels. Consistent with previous reports, Stat3^Δhep^ mice displayed notable obesity from 6 months of age, as well as fasting hyperglycemia that was observed at 12 months although not at 6 months of age (Figures S6A,B) in addition to notable hepatic steatosis at 12 months (Figures 6C). Importantly, the Stat3^Δhep^ mice also displayed significantly elevated miR-21 levels, but not miR-122, together with reduced expression of *Pparα* mRNA and a marginal reduction in its downstream target gene, *Cpt1* (Figure S6D).

### Activation of PPARα normalizes body weight, fat accumulation and glucose tolerance in mature sgp130Fc mice

To determine whether suppression of PPARα contributes to the metabolic phenotype of sgp130Fc mice, we tested the ability of the PPARα agonist, fenofibrate (Chaput et al., 2000), to rescue the sgp130Fc metabolic phenotype. We placed sgp130Fc mice and WT littermates on a normal chow diet with or without fenofibrate for 2.5 months, beginning at 4.5 months of age, prior to body weight divergence, and upon completion assessed body weight, fat accumulation, and serum glucose levels and insulin resistance (Figure 7A). Analysis of mRNA levels of PPARα target genes in the livers of naïve and fenofibrate treated sgp130Fc and WT mice confirmed the fenofibrate stimulated increase in PPARα activity. Most of the evaluated PPARα target genes in the liver, including *Cyp4a10, Cyp4a14, Fabp3, Fatp, L-Fabp* and *Vnn1* (de la Rosa Rodriguez et al., 2018; Martin et al., 1997; Montagner et al., 2016; van Diepen et al., 2014; Vida et al., 2013), were strongly elevated by fenofibrate (Figure S7); but interestingly, did not have the same effect on *Cpt-1* and *Apoe* (Figure S7). Importantly, fenofibrate treatment reduced the excess body weight (Figure 7B) and gonadal fat pad weight (Figure 7B,C) of sgp130Fc mice, bringing them to levels identical to those of fenofibrate treated WT mice, whose weight was also reduced by about 14 percent in comparison to untreated WT littermate controls. Fenofibrate treatment also reduced fasting glucose levels in the sgp130Fc mice to those of fenofibrate treated WT controls (Figure 7D), and rescued insulin sensitivity in the sgp130Fc mice (Figure 7E). Moreover, hepatic steatosis, which was varyingly present at 7 months of age in sgp130Fc mice fed a normal chow diet was strikingly reduced to WT levels by fenofibrate treatment (Figure 7F). Statistical analyses confirmed the strong strain-dependent interaction of the response to the fenofibrate diet for multiple parameters, including body weight (*P*=0.02), gonadal fat pad weight (*P*=0.005), serum glucose levels (*P*=0.008), and upregulation of *L-Fabp* mRNA expression (*P*=0.028) (Figures 7 and S7). Thus, fenofibrate treatment brought greater overall reductions in the metabolic imbalance of the sgp130Fc mice relative to the correspondingly treated WT controls, indicating that reduced PPARα levels likely constitutes part of the mechanism leading to weight gain in the sgp130Fc mice. Together, this strongly suggests that suppression of PPARα, perhaps mediated in part by miR-21 upregulation consequent to the *trans*-signaling blockade-induced reduction in Stat3 phosphorylation, contributes substantially to the aberrant metabolic phenotype of spg130Fc mice.

**Figure 7.**
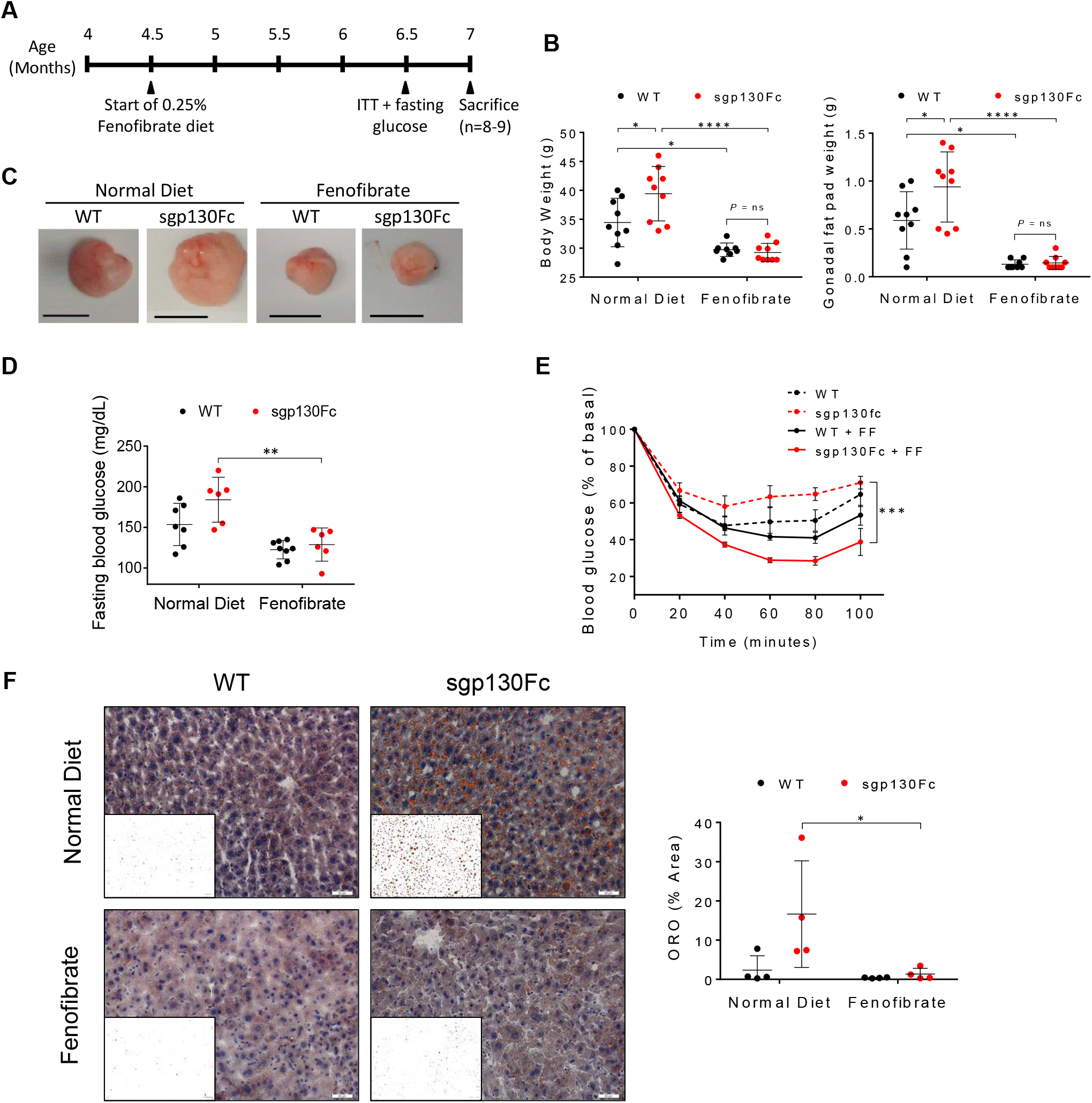
PPARα agonist fenofibrate reverses the mature-onset metabolic phenotype in sgp130Fc mice. (A) Schematic representation of the experimental design. Sgp130Fc and WT littermates aged 4.5 months were administered a Normal Chow Diet (NCD), or a NCD supplemented with Fenofibrate (FF) for 2.5 months. Fasting blood glucose levels and insulin tolerance (ITT) were assessed at 6.5 months, and the mice sacrificed at 7 months of age. (B) Body weight (left) and gonadal fat pad (right) weight in 7 months old sgp130Fc and WT littermates following NCD and FF diets (n=8-9). (C) Photographs showing representative gonadal fat pads at 7 months of age from mice fed NCD or FF diets. Scale bars, 1cm. (D) Fasting blood glucose levels in NCD or FF fed mice at the age of 6.5 months (n=6-8). (E) Insulin tolerance test (ITT) at the age of 6.5 months in sgp130Fc and WT littermates treated with NCD or FF diets (n=6-8). (F) Microphotographs showing representative ORO-stained liver sections from 7 months old NCD or FF fed mice with ImageJ processed image (inset), and quantification (right) of oil red O (ORO)-stained area in ImageJ processed images (n=4). Scale bars, 20μm. Numerical data are represented as mean ± SD (B,D,F), or mean ± SEM (F). * P < 0.05, ** P < 0.01, and *** P < 0.001 by two-way ANOVA (A,B,C,D,E,G); (F) *** P <0.001 by two-way ANOVA for comparisons of area under the curve. Significant interaction between genotype and diet were reported for: B: Body weight (\**P* = 0.02), Gonadal Fat Pad weight (\**P* = 0.05), and D (\*\**P* = 0.008).

## DISCUSSION

Co-morbidities of metabolic syndrome are an enormous healthcare burden for western societies and are becoming increasingly so for developing countries as well (Hruby & Hu, 2015). Along with weight, genetics, lifestyle, and excess caloric intake, aging is considered one of the most important independent determining risk factors in the development of metabolic syndrome (Bonomini et al., 2015). Indeed, many of the conditions contributing to metabolic syndrome, such as obesity, insulin resistance, and inflammation, also increase in prevalence during aging (Bonomini et al., 2015). IL-6 has long been seen as a pro-inflammatory cytokine, with functions that parallel those of TNFα as a driver of obesity-associated metabolic complications. However, ample evidence also indicates that IL-6 signaling is a crucial regulator of metabolic homeostasis, and, importantly, contributes centrally through a mechanism of *trans*-signaling to affect body weight control, reduce feeding, and maintain peripheral glucose tolerance, especially in obesity (Timper et al., 2017).

Our findings demonstrate that in aging *peripheral trans*-signaling also contributes to the control of behavioral determinants of energy expenditure and to maintain metabolic homeostasis. Thus, following *tran*s-signaling blockade energy expenditure imbalance is manifest at an early age with hyperphagia balanced by hyperactivity and in aging with ectopic lipid accumulation, including hypertrophy-related adiposity and hepatic steatosis, glucose intolerance, and insulin resistance, together with reduced physical activity.

Blockade of peripheral *trans*-signaling in sgp130Fc mice recapitulates many of the metabolism-related phenotypic aspects found in some strains of IL-6^-/-^ and also Stat3^ΔHep^ mice (Inoue et al., 2004; Matthews et al., 2010; Wallenius et al., 2002), but with striking and perhaps instructive differences. For instance, sgp130Fc mice display hyperphagia from an early age, while obese IL-6^-/-^ mice and Stat3^ΔHep^ mice do not. While the molecular and cellular mechanism(s) leading to hyperactivity and hyperphagia in sgp130Fc mice are not entirely clear, our observation that ectopic sgp130 expression reduces leptin signaling in the hypothalamus is in agreement with previous reports that IL-6 *trans*-signaling in the hypothalamus functions to limit food-intake especially in obesity (Timper et al., 2017). However, in our model, the sgp130Fc transgene is expressed peripherally under the PEPCK promoter, and the recombinant protein does not cross the blood-brain barrier (Braun et al., 2013; Rabe et al., 2008).

How peripheral *trans*-signaling blockade reduces leptin signaling in the hypothalamus is presently unclear. Various mechanisms have been shown to underlie leptin resistance in obesity in experimental models. These include reduced leptin passage across the blood-brain barrier into the CNS, which can also be inhibited by triglycerides and TNF-α, impairment of leptin receptor (LepRb) trafficking, suppression of LepRb signaling, and also environmental effects such as neonatal overfeeding (Banks et al., 2004; Cui et al., 2017; Jung & Kim, 2013). Thus, it is conceivable that t*rans*-signaling blockade in the periphery may impede leptin signaling indirectly by affecting one of these pathways. Alternatively, sgp130Fc may act directly to impede leptin signaling. Further studies are required to elucidate the mechanism(s) through which peripheral *trans*-signaling contributes to mediate the cross-talk between the periphery and CNS regulated behavioral functions of appetite and activity.

The observation that young spg130Fc mice do not display weight gain despite their hyperphagia suggests that factors distinct from hyperphagia may also contribute to obesity and metabolic imbalance also in these mice. Glucose intolerance in IL-6 signaling-deficient mice challenged with an HFD has consistently been found to be associated with an exacerbated inflammatory response in the liver and adipose tissue, suggesting that IL-6 may have a unique homeostatic role in limiting the low-grade inflammatory state thought to be a critical driver in the development of glucose intolerance (Matthews et al., 2010; Mauer et al., 2014; Wunderlich et al., 2010). However, in our sgp130Fc mice, *tran*s-signaling blockade induced glucose intolerance in the absence of hepatic inflammation. This observation suggests that *trans*-signaling mediated prevention of glucose intolerance and its role in metabolic homeostasis may operate via a mechanism(s) other than the limitation of low-grade hepatic inflammation.

Here we identify PPARα suppression in the liver to be a critical mechanism in the onset of mature-onset weight gain, hepatic steatosis, and insulin resistance in *trans-*signaling deficient mice, which may be mediated by reduced STAT3 suppression of miR-21 expression. This conclusion is consistent with previous reports showing PPARα to be an important, miR-21-targeted regulator of metabolism, which is also reduced in human NAFLD patients (Liss & Finck, 2017), and with reports demonstrating that miR-21 is suppressed by STAT3 (Zhang et al., 2011). However, considering that the phenotypes associated with *trans*-signaling blockade are broader in nature and appear earlier than those associated with hepatocyte-targeted STAT3 ablation alone, it is also clear that suppression of STAT3 and PPARα in hepatocytes can account only in part for the phenotype induced by *trans*-signaling blockade. Curiously, while PPARα suppression in sgp130Fc mice was evident from an early age, the metabolic consequences of IL-6/STAT3 signaling-disruption described here and by others (Inoue et al., 2004; Matthews et al., 2010; Wallenius et al., 2002) are strongly age-dependent, appearing in mice only after about 6 months of age. This discrepancy suggests that critical factors may come into play in aging to allow the metabolic imbalance induced by *trans*-signaling blockade and PPARα suppression. In this sense, the reduction in physical activity displayed by mature sgp130Fc mice, which is reminiscent of the sedentary behavior displayed by some adult humans, may be of relevance; although, how *trans*-signaling interacts with aging-related factors to modify physical activity is presently unclear.

It is perhaps important to note that in addition to IL-6 *trans*-signaling, sgp130 has also been observed to partially inhibit signaling by other IL-6 family members *in vitro*, including, interleukin-11 leukemia inhibitory factor (LIF), ciliary neurotrophic factor, and oncostatin M (OSM), albeit with about 100-fold lower efficacy (Jostock et al., 2001; Narazaki et al., 1993). Thus, it is reasonable to expect that the metabolic phenotype observed in sgp130Fc mice may be broader in scope than that observed in IL-6 *knockout* mice, and is consistent with the broad roles of IL-6 family cytokines in metabolic homeostasis (Pasquin et al., 2016).

Lastly, it is intriguing that although these aberrant metabolic traits appear common to many strains of mice that carry mutations affecting IL-6/STAT3 signaling, they reportedly are not shared by all strains of IL-6^-/-^ and Stat3^Δhep^ mice, despite carrying similar or identical mutant alleles (Di Gregorio et al., 2004; Moh et al., 2007). In line with this observation, our IL-6^-/-^ mice displayed mature-onset hepatic steatosis, but do not become notably obese. Similarly, the obese phenotype of aged sgp130Fc mice seen here has not been observed in other globally dispersed sgp130Fc colonies. These include the sgp130Fc colonies located in Hamburg and Kiel, which did not show the herein described obese phenotype (SRJ, personal communication); although, a trend of increasing fat mass has been observed by others in young sgp130Fc mice (Kammoun et al., 2017). These observations suggest that subtle, and yet unidentified genetic, epigenetic, or perhaps environmental factors may influence the penetrance of metabolic phenotypes related to IL-6, STAT3, and *trans*-signaling deficiencies. By analyzing the microbiome of sgp130Fc and WT strains maintained in animal facilities in three different geographical locations (Jerusalem, Israel, and Kiel and Hamburg, Germany), we find major differences between the animal houses, which by PCoA analysis are highly statistically significant (*P* = 0.001) (Figure S8, Table SI). The microbiome in Jerusalem has a higher Clostridial population and fewer Bactericides. The literature perceives Bacteroides as a beneficial Genus and Clostridial as a harming Genus (Wexler, 2007; Woting et al., 2014). The metabolic effects we see in the present study, although being strain-dependent are possibly also affected by the differences in the microbiome seen in the different animal facilities. These observations warrant further investigations into the connection between specific bacterial strains and their metabolic effects.

Taken together, our findings demonstrate a role for peripheral IL-6 *trans*-signaling in metabolic homeostasis in aging and thus provide clinical significance to the apparent association of elevated sgp130 levels with obesity, insulin resistance, and diabetes found in some human patients (Nikolajuk et al., 2010; Weiss et al., 2013; Wonnerth et al., 2014; Zuliani et al., 2010). Future research is required to unravel the mechanism(s) through which peripheral *tran*s-signaling affects leptin resistance, food-intake and activity, and to identify aging-related determinants that it appears to counter in maintaining metabolic homeostasis. Lastly, our experiments identify sgp130 and IL-6 *trans-*signaling affected pathways as potential drug targets for obesity and obesity-associated insulin and leptin resistance in aging.

## Materials and Methods

### Animal Care

Mice were maintained in an animal facility with a temperature of ∼23°C in a 12-hour light-dark cycle, under SPF conditions and received sterile commercial rodent chow and water *ad libitum*. Maintenance of mice and all experimental procedures were performed in accordance with the Institutional Animal Care and Use Committee approved animal treatment protocols (license number OPRR-A01-5011).

### Genetic mouse models

Male mice were used in all experiments. Sgp130Fc^+/+^ (C57BL/6N) mice (Rabe et al., 2008) were crossed with WT (C57BL/6J) mice purchased from Harlan Laboratories (Jerusalem, Israel) to generate heterozygous sgp130Fc^+/-^ mice. IL-6^+/-^ mice were then crossed in order to generate both homozygous IL-6^-/-^ mice and wild type (WT) (IL-6^+/+^) littermates. Stat3^flox/flox^ Alb-Cre/ (Stat3^ΔHep^) mice were generated from STAT3 (Stat3^flox/flox^) (C57BL/6) mice crossed with Alb-Cre^+/+^ (C57BL/6) mice. See Supplementary Information for further details in the generation and validation of genetic mouse models.

### Study design

Male mice were used in all experimental groups were housed in groups consisting of 2-3 mice per cage. Sgp130Fc mice and WT littermates were raised to the age of 14 months. Experimental cohorts were sacrificed at 2, 5, and 14 months of age. Stat3^ΔHep^ and Stat3^flox^ littermates were grown to 12 months of age. Upon sacrifice of the cohorts, liver and adipose tissue samples were removed and snap-frozen in liquid nitrogen for protein and RNA extraction, or embedded for frozen tissue sections or fixed in 4 percent formaldehyde and paraffin-embedded for histological and immunostaining analyses. For full details analyses concerning MRI, metabolic cages, food intake, leptin resistance, glucose and insulin tolerance testing, insulin secretion, insulin signaling, Western blot, histochemical and immunostaining, RNA, flow cytometry, and taxonomic microbiota see Supplementary Information.

### Statistical Analysis

Data were evaluated for significance by two-tailed Student’s *t*-test or Mann-Whitney test, unless otherwise noted. *P* ≤ 0.05 was considered significant for all analyses. Calculations were performed using GraphPad Prism 6.02 software (Graph-Pad Software, Inc, San Diego, CA) unless otherwise noted.

## Acknowledgments

The authors are grateful to Deborah Olam for excellent technical assistance; to Catherine Tempel-Brami and Yael S. Schiffenbauer for assistance in performing MRI experiments; to Neta Barashi, Ann Reisch Saada for helpful discussions; and to Tali Bdolah-Abram for expert advice concerning statistical evaluations.

## Supplemental Figures

**Figure S1. (Related to Figures 1 and 2).**
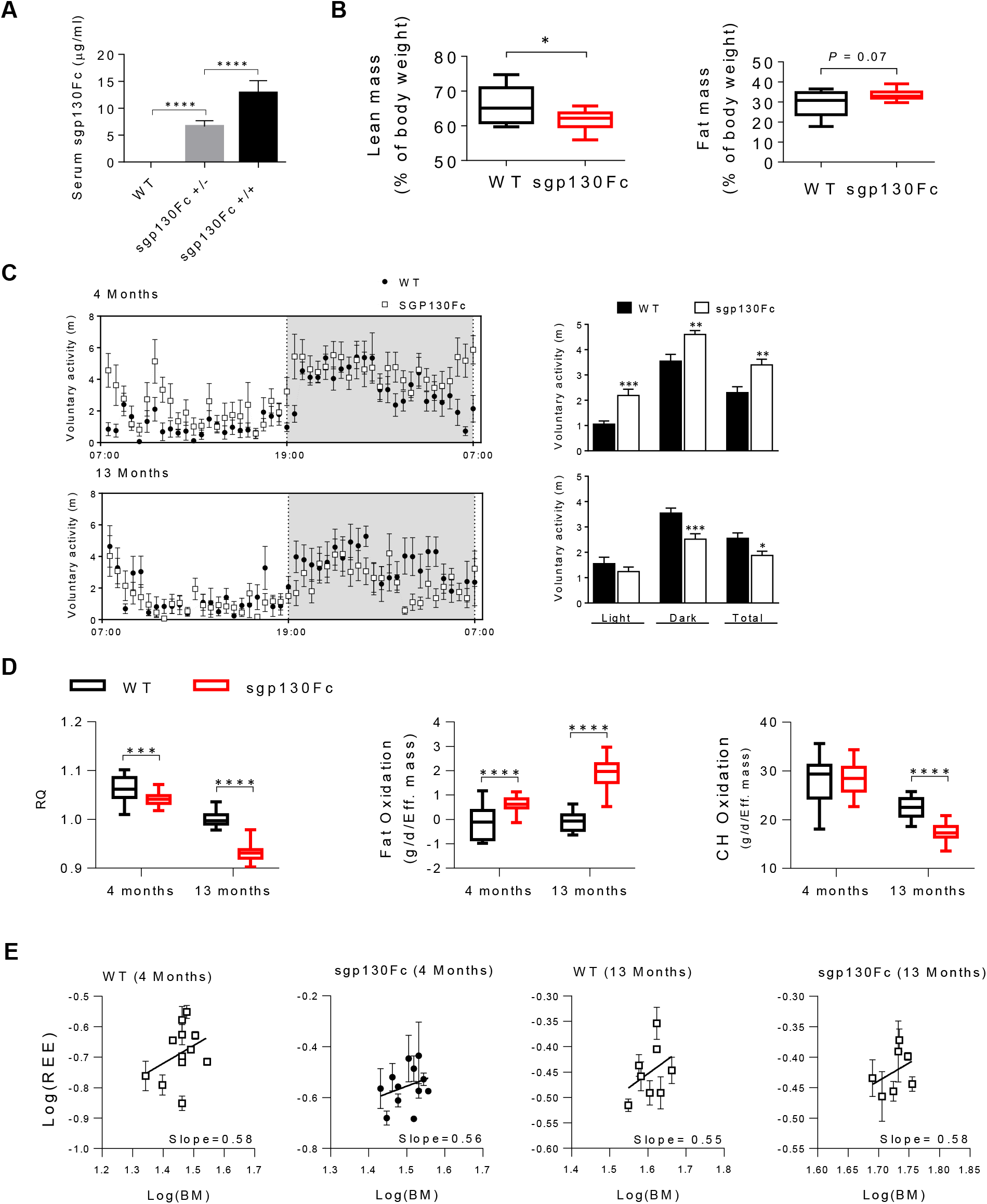
IL-6 *trans*-signaling blockade increases fat mass and decreases lean mass associated with decreased energy expenditure. (A) Serum concentrations of human sgp130Fc by ELISA in wild type (WT), heterozygous (-/+) and homozygous (+/+) sgp130Fc mice at 2 months of age (n=8-15). (B) Echo MRI analysis of lean and fat body mass as a function of % body weight in sgp130Fc and WT littermates at 14 months of age (n=9-10). (C) Voluntary activity (pedmeter) at 4 (n=12) and 13 (n=8) months of age. (D) Metabolic activity for dark cycle including respiratory quotient (RQ), fat oxidation and carbohydrate (CH) oxidation of WT and sgp130Fc mice at 4 (n=12) and 13 months (n=8). (E) Allometric regression analysis for determination of mass scaling exponent of WT and sgp130Fc mice. Data are represented as mean ± SD (A,C), or box and whiskers plots (B,D) \**P* < 0.05, \*\**P*<0.01, ** *P* <0.01 \*\*\**P*<0.001, and \*\*\*\**P*<0.0001 by two-tailed, Student’s *t* test (A-C), or by two-way ANOVA (D). Significant interactions (\*\*\*\**P* < 0.0001) between genotype and age were returned for RQ, fat oxidation and carbohydrate oxidation.

**Figure S2. (Related to Figures 3 and 5).**
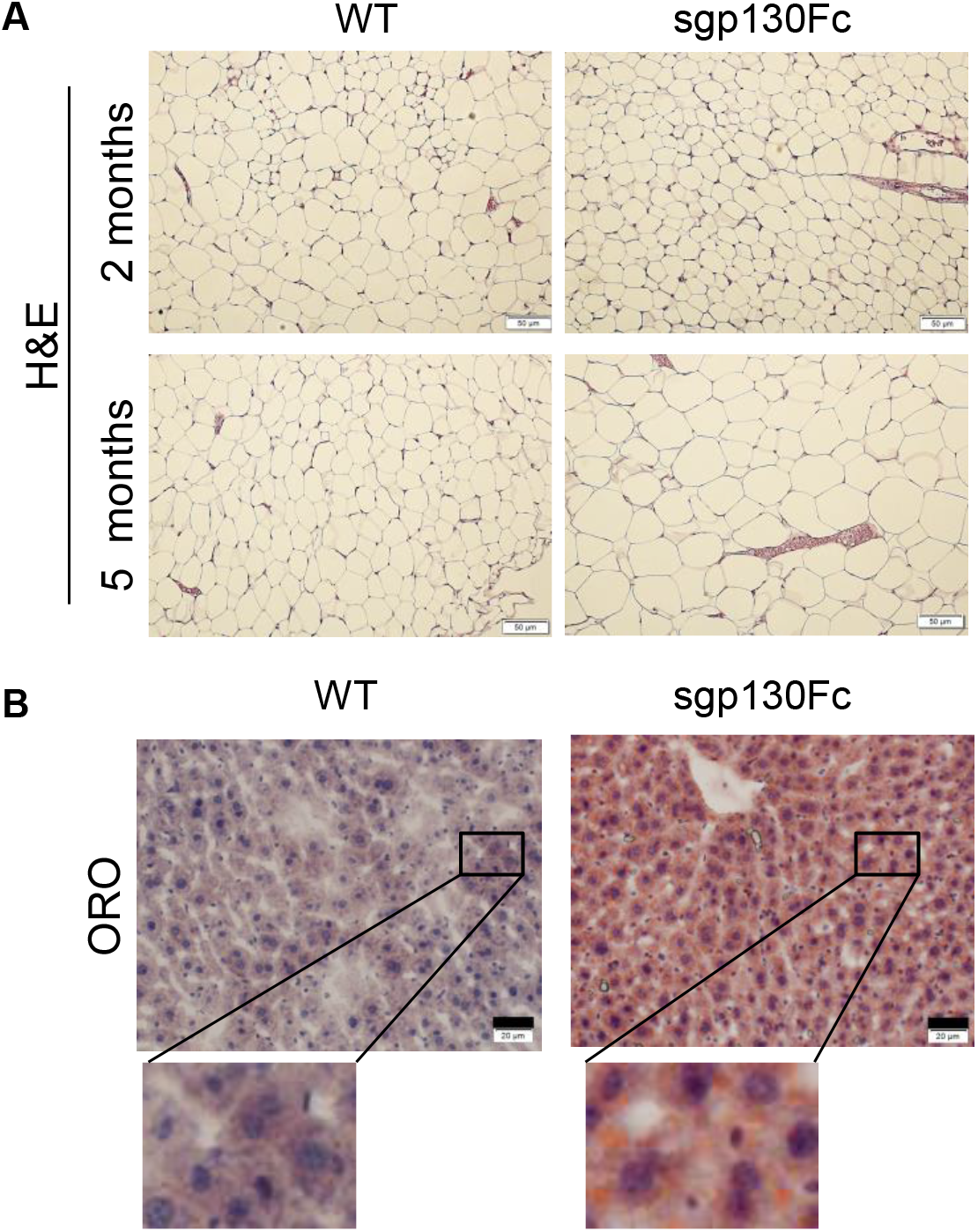
Inhibition of IL6 *trans*-signaling results in adipocyte hypertrophy and steatosis at 5 months of age. (A) H&E stained sections from paraffin embedded adipose tissue from sgp130Fc and WT littermates at 2 and 5 months of age. Scale bars, 50μm. (B) ORO-stained liver sections from sgp130Fc and WT littermates at the age of 5months. Scale bars, 20 μm.

**Figure S3. (Related to Figure 4).**
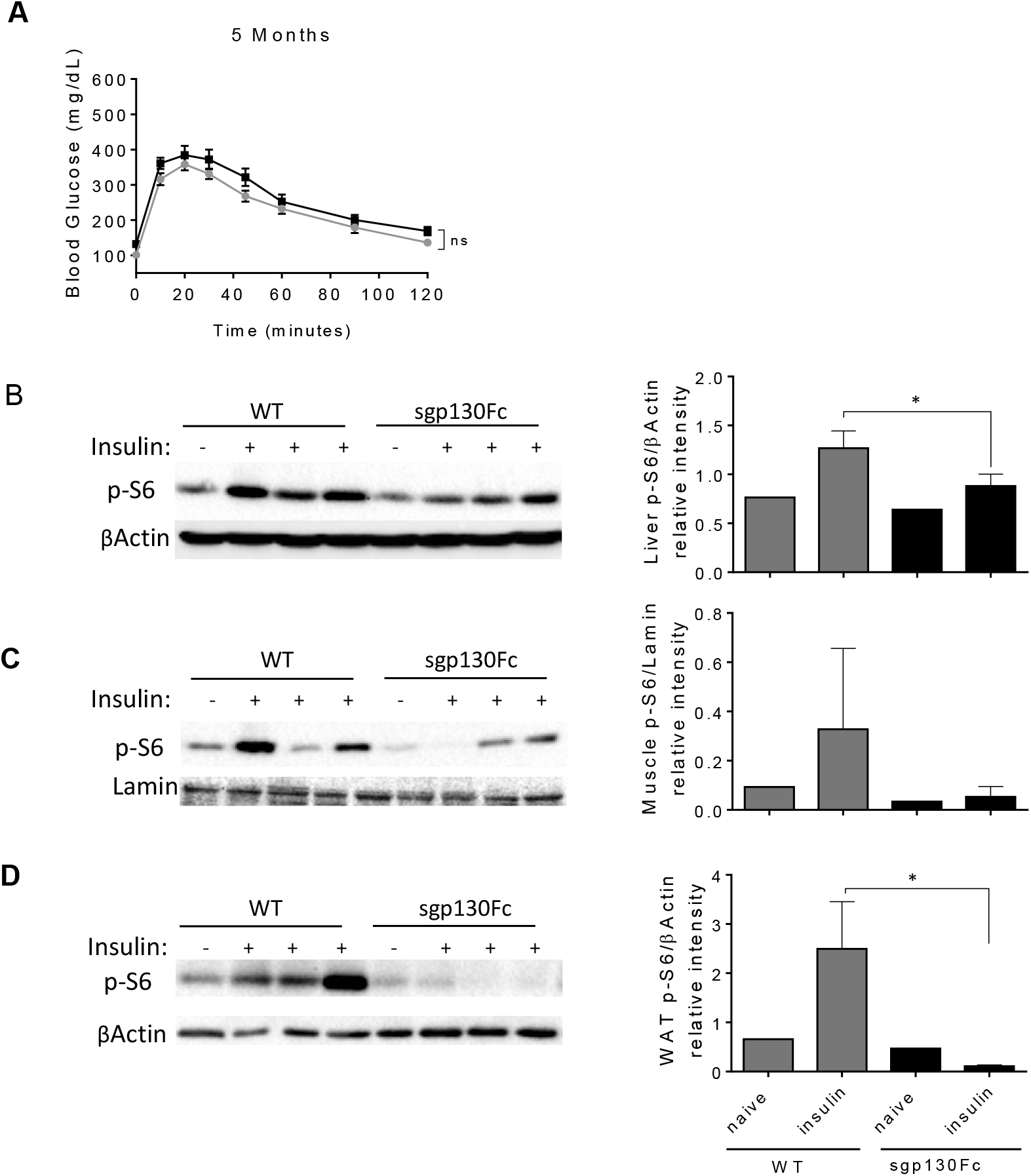
Inhibition of IL-6 *trans*-signaling induces glucose intolerance and peripheral insulin resistance. (A) Glucose levels before and after glucose challenge in sgp130Fc versus WT mice at 5 (left) months of age (n=7). (B-D) Phospho-S6 (pS6) protein levels by Western Blot analysis of liver (B), muscle (C), and white adipose tissue (WAT) (D) tissues before and 10 minutes post insulin injection in 10 month-old sgp130Fc and WT mice. Quantification of band intensity by ImageJ (right). (n=3 for groups of insulin-treated mice). Data are represented as mean ± SEM (A), or mean ± SD (B-D). (A) *P = ns* for comparisons of area under the curve by two-tailed Student’s *t* test. (B, D) * *P* < 0.05, by two-tailed Student’s *t* test.

**Figure S4. (Related to Figures 1,2,3 and 5).**
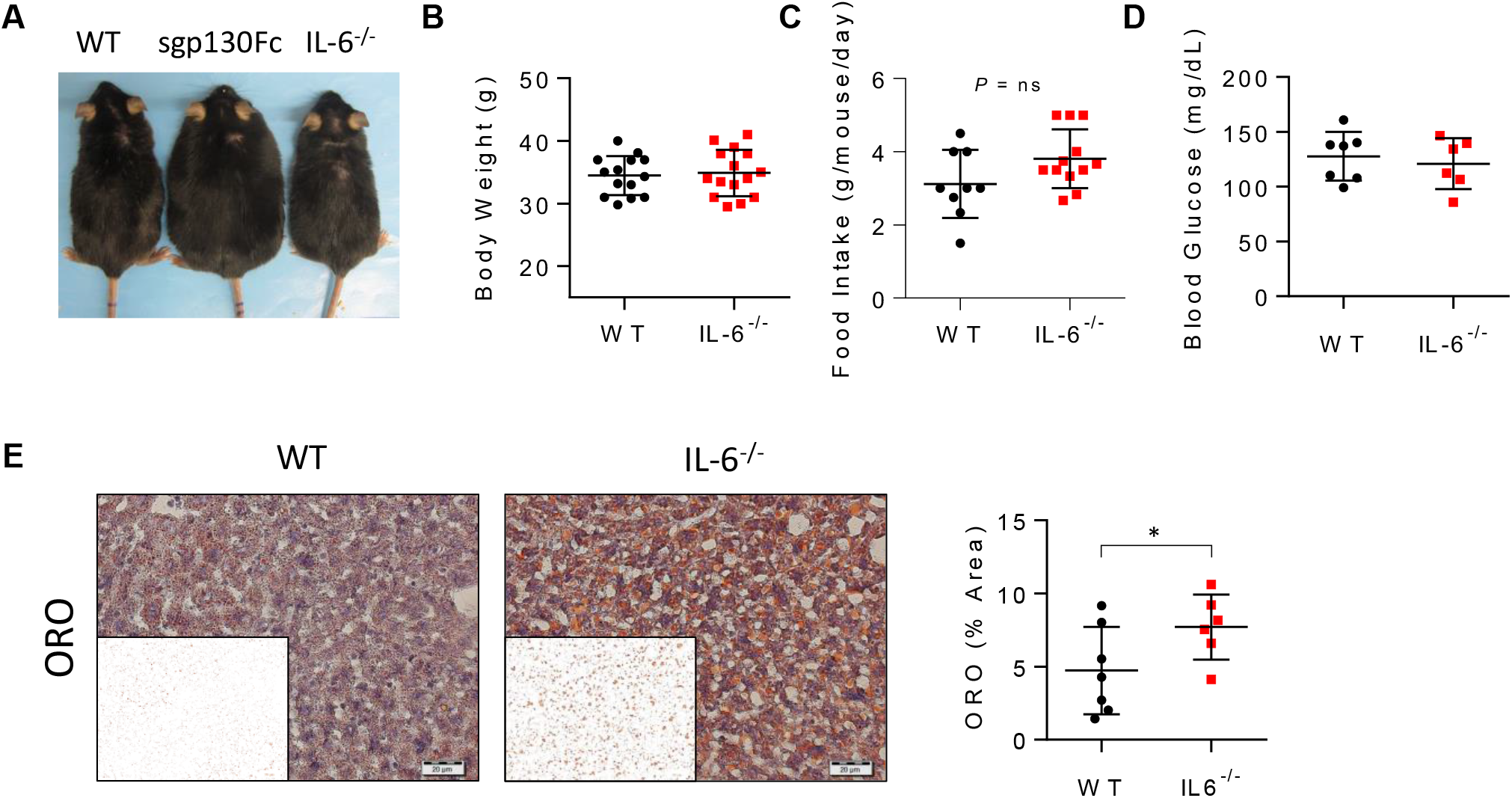
IL-6 *knockout* mice display hepatic steatosis, but not obesity. (A) Representative photographs of WT,sgp130Fc, and IL-6^-/-^ mice at the age of 12 months. (B) Body weight at 8 months of age in WT and IL-6^-/-^ littermates (n=14). (C) Daily food-intake at 6 months of age (n=9-12). Data represent mean food intake per mouse/cage per day averaged over a 2 week period. (D) Fasting glucose levels at 6 months of age in WT and IL-6^-/-^ littermates, (n=6-7). (E) Representative microphotographs showing steatosis by oil red O (ORO) staining in liver sections from WT and IL-6^-/-^ littermates at 8 months of age and ImageJ processed image (Inset). Scale bars represent 20 μm. Quantification of ORO-stained area by ImageJ analysis (right) (n=6-7). Data are represented as mean ± SD. * *P* < 0.05 by two-tailed, Student’s *t* test.

**Figure S5. (Related to Figures 5).**
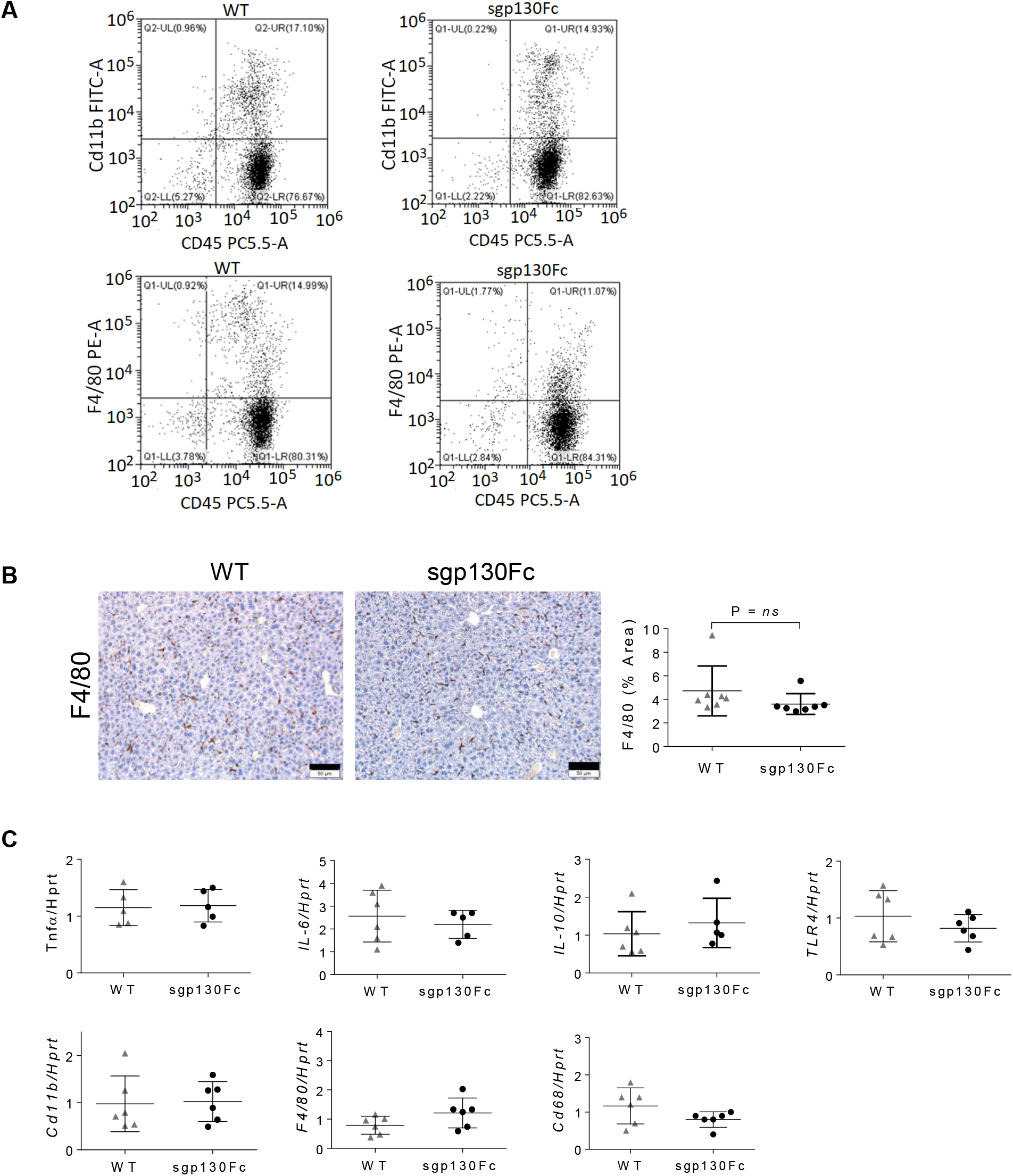
Inhibition of *trans*-signaling induces steatosis without hepatic inflammation. (A) Representative plots from flow cytometry of double-positive Cd45+/Cd11b+ and Cd45+/F4/80+ cells (upper right quandrants) in livers of sgp130Fc and WT littermates at 11 months of age. (B) Representative photomicrographs of F4/80 immunostained paraffin embedded liver sections from 5-month old sgp130Fc and WT littermates and quantification (right) of F4/80+ cells as a function of area stained by ImageJ analysis, (n=8). Scale bars, 50μm. (C) Hepatic mRNA expression levels by real time qPCR of indicated inflammatory and monocyte markers in sgp130Fc vs. WT littermates at the age of 14 months. Data are represented as mean ± SD. *P < 0.05 by two-tailed, Student’s *t* test.

**Figure S6. (Related to Figure 6).**
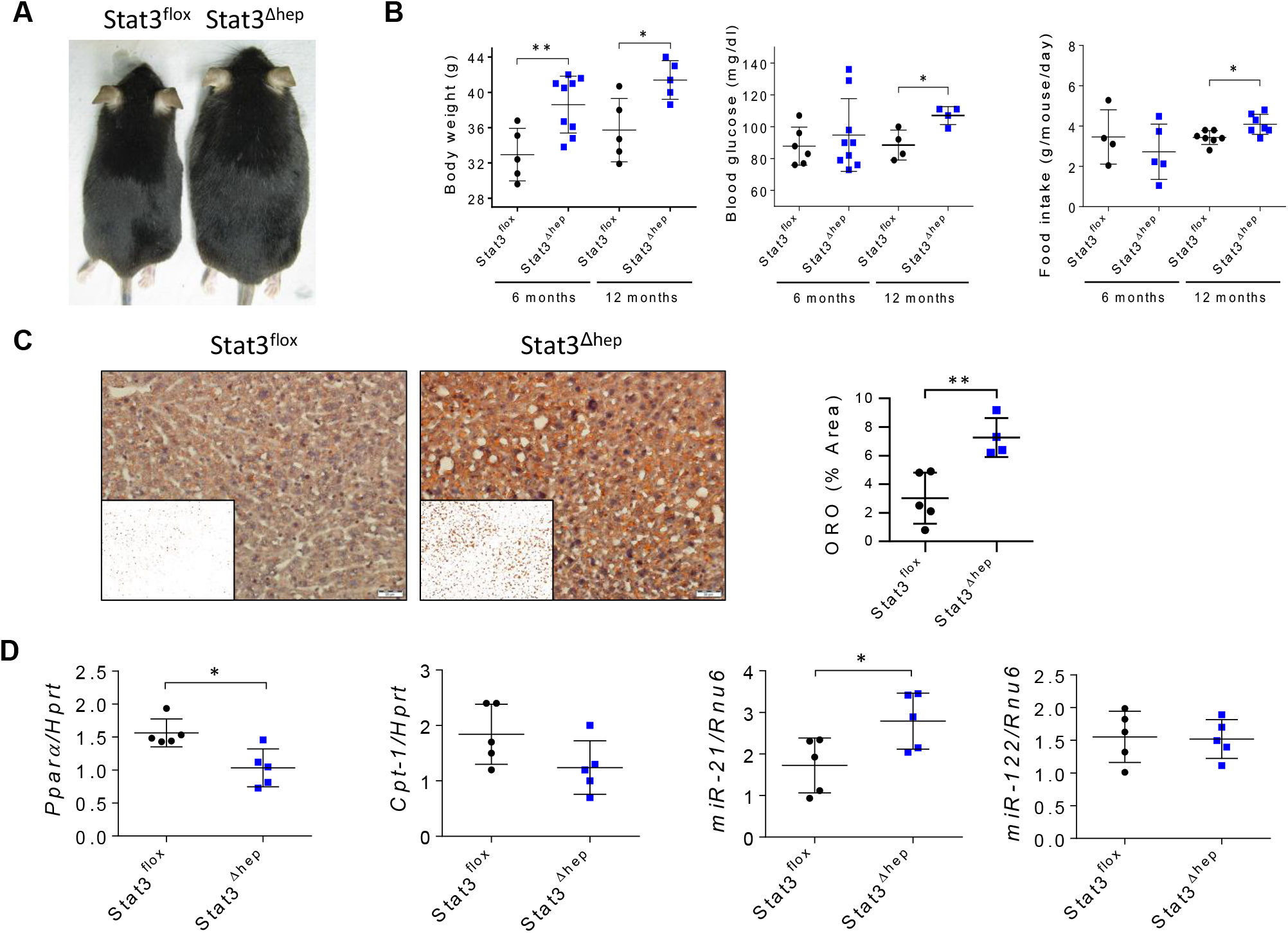
Trans-signaling protection against mature-onset metabolic syndrome may be mediated by Stat3 in hepatocytes. (A) Photographic image showing representative Stat3^Δhep^ and Stat3^floxP^ mice at 12 months of age. (B) Body weight, fasting blood glucose levels, daily food intake, and fasting-induced weight loss in 12 month old Stat3^Δhep^ and Stat3^floxP^ mice, (n=5-8). (C) Representative oil red O (ORO) stained liver sections from Stat3^Δhep^ and Stat3^floxP^ mice at 12 months of age with ImageJ processed image (inset), and quantification of ORO-stained areas by ImageJ (right), (n=4-5). Scale bars, 20μm. (D) Real time qPCR quantification of mRNAs encoding PPARα, Cpt-1, and miR-21, and miR-122 in livers of Stat3Δhep and Stat3floxP mice at 12 months of age (n=4-5). Data are represented as mean ± SD. * P < 0.05, and ** P < 0.01 by two-tailed, Student’s t test.

**Figure S7. (Related to Figure 6).**
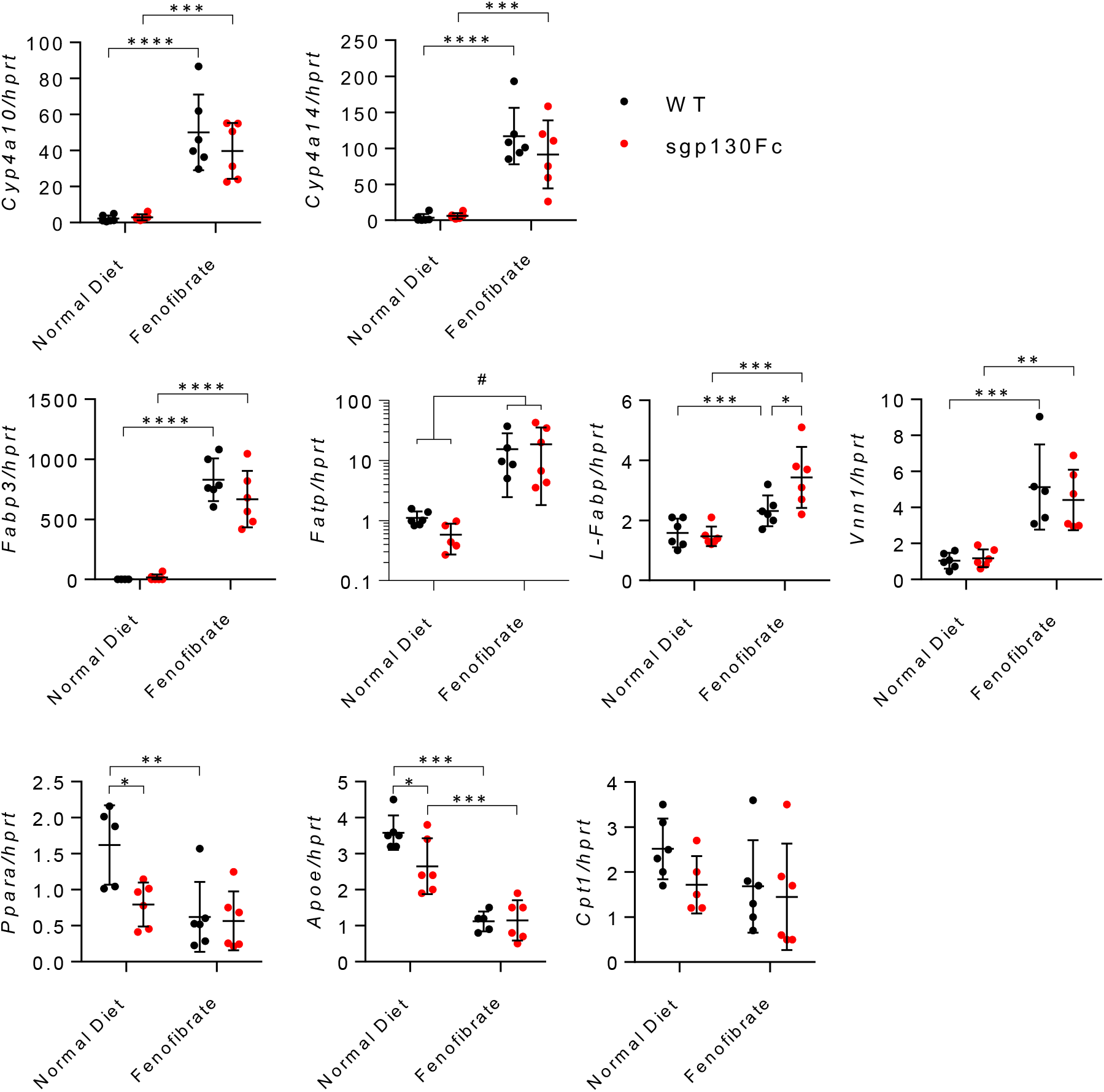
The effect of a fenofibrate on *PPARα* and downstream target gene expression in the liver. Sgp130Fc and wild type (WT) littermates aged 4.5 months were administered a Normal Chow Diet (NCD), or a NCD supplemented with Fenofibrate (FF) for 2.5 months. Data show hepatic mRNA levels of PPARα and down-stream *PPARα* target genes, *Apoe, Cpt1, Cyp4a10, Cyp4a14, Fabp3, Fatp, L-Fabp* and *Vnn1* by real time qPCR analysis, (n=6). Data are represented as mean ± SD. * *P*<0.05, ** *P*<0.01, *** *P*<0.001 by two-way ANOVA. # denotes main effect (*P*<0.01) for diet by two-way ANOVA. Significant interaction between genotype and diet were reported for *Fatp* (*P*=0.028).

**Figure S8.**
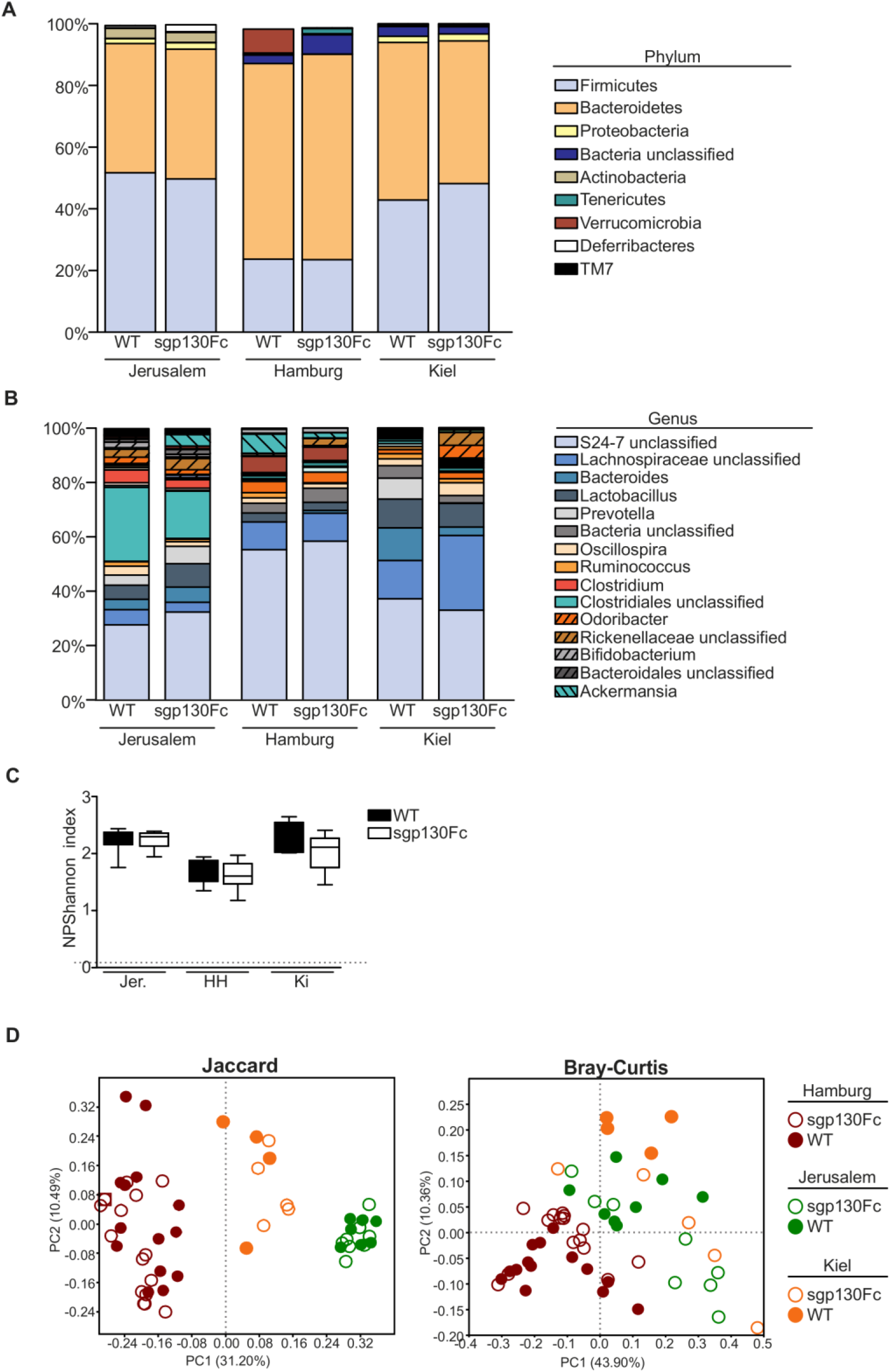
Inhibition of IL-6 trans-signalling does not alter gut microbiota composition. **(A)** Gut microbiota composition at the phylum level based on 16S rRNA gene sequencing in feces. **(B)** Relative bacterial abundance at genus level does not show alterations that can be associated with the genotype in three independent animal facilities. **(C)** Box plots of the Shannon diversity Indices show that total gut microbial diversity is independent of mouse genotype in three independent animal facilities. **(D)** *left panel:* Principal coordinate analysis (PCoA) on Jaccard distance matrices demonstrates alterations in microbiota composition between different animal facilities. *right panel:* PCoA based on Bray-Curtis dissimilarity index of bacterial genus abundances.

**Table SI:** *P*-values are shown for permutational analysis of variance (PERMANOVA) with Bonferroni correction for multiple testing

|  | *Jaccard* |  |  | *Bray-Curtis* |  |  |
| --- | --- | --- | --- | --- | --- | --- |
| **facility** | Jerusalem | Hamburg | Kiel | Jerusalem | Hamburg | Kiel |
| Jerusalem |  | 0.0003 | 0.0003 |  | 0.0003 | 0.0942 |
| Hamburg | 0.0003 |  | 0.0003 | 0.0003 |  | 0.0003 |
| Kiel | 0.0003 | 0.0003 |  | 0.0942 | 0.0003 |  |

