## Supplemental Methods for "Peripheral sgp130-mediated *trans*-signaling blockade induces obesity and insulin resistance in mice via PPARα suppression"

### SUPPLEMENTAL INFORMATION

#### Supplemental Materials and Methods

##### Genetic mouse models

Male mice were used in all experiments. Sgp130Fc$^{+/+}$ (C57BL/6N) mice (Rabe et al., 2008) were crossed with WT (C57BL/6J) mice purchased from Harlan Laboratories (Jerusalem, Israel) to generate heterozygous sgp130Fc$^{+/-}$ mice. Heterozygous sgp130Fc$^{+/-}$ mice were then crossed in order to generate both homozygous sgp130Fc$^{+/+}$ mice and wild type (WT) (sgp130Fc$^{-/-}$) littermates. Homozygosity of the sgp130Fc$^{+/+}$ and sgp130Fc$^{-/-}$ (WT) alleles was determined by ELISA for soluble sg130Fc protein levels in the serum using anti-human sgp130 ELISA (R&D, cat.no. DY228) (Figure S1A). IL-6$^{-/-}$ (C57BL/6) mice were purchased from The Jackson Laboratory (Bar Harbor, ME) and crossed with WT (C57BL/6J) mice to form heterozygous IL-6$^{+/-}$ mice. IL-6$^{+/-}$ mice were then crossed in order to generate both homozygous IL-6$^{-/-}$ mice and wild type (WT) (IL-6$^{+/+}$) littermates. Floxed-STAT3 (Stat3$^{flox/flox}$) (C57BL/6) mice (Takeda et al., 1998) [kindly provided by E. Razin (Hebrew Univ. of Jerusalem, Israel)] were crossed with Alb-Cre$^{+/+}$ (C57BL/6) mice (Postic et al., 1999) [kindly provided by D. Wallach (Weizmann Inst. of Science, Rehovot, Israel)] to form Stat3$^{flox/+}$ Alb-Cre$^{+/-}$ mice, the offspring of which were then crossed to generate Stat3$^{flox/flox}$ Alb-Cre/ (Stat3$^{\Delta Hep}$) and Stat3$^{flox}$ strains. Genotyping analysis was performed by PCR analysis using the following DNA primer sets: STAT3$^{flox}$ (sense) 5'-cctgaagaccaagttcatctgtgtgac and (antisense) 5'-cacacaagccatcaaactctggtctcc; Stat3$^{\Delta Hep}$: (sense)5'-gatttgagtcagggatccataacttcg and (antisense) 5'-cacacaagccatcaaactctggtctcc; Alb-Cre: (sense) 5'-tatcttctatatcttcaggcgc and (antisense) 5'- gtgaacgaacctggtcgaaatcag.

##### MRI Analysis

*In vivo* MRI analysis was performed using the M2 (Aspect Ltd, Israel), a compact, high-performance MRI system, equipped with a 35-mm mouse whole-body coil (Tempel-Brami et al., 2015). For *in vivo* MRI imaging, mice were maintained in an anesthetized state with 2% isoflurane and placed on a specially designed heated bed where physiological signals were monitored throughout the experiment to ensure the animals' wellbeing. MRI acquisition parameters include spin echo with slice thickness = 1 mm, repetition time = 450 ms, echo time = 11.5 ms, field of view = 70 mm, matrix = 256 X 256, and acquisition time = 3.5 min. Fat content was analyzed using Analyze 7.0 (Analyze Direct, USA) software.

#### Metabolic cage analysis

Mice were metabolically assessed by using the Promethion High-Definition Behavioral Phenotyping System (Sable Instruments, Inc., Las Vegas, NV, USA) as described previously (Udi et al., 2017). Briefly, mice with free access to food and water were subjected to a standard 12 h light/12 h dark cycle, which consisted of a 48 h acclimation period followed by 24 h of sampling. Respiratory gases were measured by using the GA-3 gas analyzer using a pull-mode, negative-pressure system. Airflow was measured and controlled by FR-8, with a set flow rate of 2000 mL/min. Water vapor was continuously measured and its dilution effect on O2 and CO2 was mathematically compensated. Total Energy Expenditure (TEE) was calculated according to the method of Tschöp et al 2011 (Tschop et al., 2011). As such, instead of dividing the metabolic rate by Kg effective mass to the power of 0.75 (Kleiber's exponent), we have performed an ANCOVA analysis and used this value to recalculate the TEE. Briefly, we took the logarithms of body mass and of the resting metabolic rate and regressed them against each other (Figure S1E). The slopes of the regression were the power to which body mass was raised in order to yield the panels describing TEE, in proportion to their metabolic rate. Ambulatory activity and position were monitored simultaneously with the collection of the calorimetry data using XYZ beam arrays with a beam spacing of 0.25 cm.

#### Food intake

Food intake was assessed by measurement of pre-weighed food pellets remaining after a 24-hour period. The average daily food-intake per mouse was defined as the average food consumption per day per cage measured over a two-week assessment period and divided by the number of mice per cage.

#### Leptin resistance

Following an overnight fast, sgp130Fc and WT littermates were injected with recombinant murine leptin (3mg/kg, i.p) (Peprotech #450-31) or carrier control (0.1%BSA in PBS). Hypothalamus tissue was removed by dissection 45 minutes’ post injection, snap frozen, and stored at -80°C until protein extraction and analysis by Western blot.

#### Glucose and insulin tolerance tests

Glucose tolerance (GTT) and insulin tolerance (ITT) tests were performed 2 weeks apart on fasted mice. For GTT analysis, mice received an intraperitoneal (IP) injection of dextrose (Merck) at a dosage of 1.5g/kg. Glucose levels were measured at 10, 20, 30, 40, 60, 90, and 120 minutes post-dextrose injection. ITT analysis was performed in mice by IP injection of human insulin (Actrapid) at a dosage of 0.87units/kg followed by the assessment of blood glucose levels every 20 minutes for 2 hours. Blood glucose levels were assessed using an Accu-Chek® blood glucometer and glucose test strips.

##### Insulin secretion

Insulin was measured in plasma samples collected from mice following overnight fasting, and 15 minutes following administration by gavage of a liquid meal consisting of Ensure Plus® supplemented with 24% (w/v) dextrose at a dextrose dosage of 2g/kg body weight. Blood samples were collected in EDTA-coated tubes (MiniCollect, Greiner Bio-one) containing protease inhibitor cocktail (Calbiochem) according to the manufacturer's instructions. Plasma samples were stored at -80º C and analyzed for insulin by Multiplex ELISA (Milliplex mouse metabolic bead panel, Millipore).

##### Analysis of insulin signaling in peripheral tissues

Sgp130Fc and WT littermates were fasted overnight and then sacrificed 10 minutes following injection of saline or human insulin (Actrapid) (1unit/kg, i.p.). Liver, muscle, and adipose tissue samples were snap-frozen in liquid nitrogen and stored at -80º C for protein extraction and western blot analysis.

##### Western blot analysis

Protein extracts were prepared from frozen tissue (~50 mg) by homogenization in lysis buffer (1% NP-40, 10 mM Tris pH 7.8, 150 mM NaCl, 40 mM EDTA, 10 mM Na-Pyrophosphate, 10 mM NaF, 1mM PMSF, 4 mM Orthovanadate, Mini complete iprotease inhibitor®), separated by polyacrylamide gel electrophoresis and subjected to Western blot analysis. For analysis of phosphorylated ribosomal protein S6 (p-S6), western blots were probed with an antibody against p-S6 ribosomal protein (Ser240/244, Cell Signaling) followed by HRP-conjugated anti-rabbit antibody (Dako). For analysis of PPARα, blots were probed with anti-mouse PPARα (Abcam) followed by HRP-conjugated anti-rabbit antibody (Dako). For analysis of phosphorylated STAT3, blots were probed with anti-phosphorylated STAT3 (Santa Cruz), followed by HRP-conjugated anti-mouse antibody (Dako). Western blots were developed with EZ-ECL kit (Biological Industries). Blots were stripped with 0.1 M glycine pH 2.2 and re-probed with a mouse anti-β-actin antibody (Sigma) as a loading control for hypothalamus, liver and adipose tissues or with a rabbit anti-Lamin B1 antibody (Novusbio) as a loading control for muscle tissue. Quantification of band intensities was performed using ImageJ software.

#### Histology and Immunohistochemical staining

Livers and adipose samples were placed in 4% buffered formaldehyde for 24 hours, followed by 80% ethanol and then embedded in paraffin blocks. Liver and adipose tissue sections (5 µm) were deparaffinized with xylene, and hydrated through graded ethanol and stained for H&E by standard procedures. Adipocyte cell size was quantified from H&E stained thin sections using ImageJ software. Macrophages were stained using rat anti-mouse F4/80 antigen (Serotec), followed by anti-Rat HRP (Histofine) and developed with a DAB kit (Zymed). Oil red O (ORO) (Sigma) staining was performed on liver frozen sections (10µm) fixed in 0.5% Glutaraldehyde (Sigma) and counterstained with hematoxylin (Emmonya Biotech). Images of stained sections were quantified as percentage area stained positively per high power field was quantified using ImageJ software (ImageJ, RRID: SCR_003070) in 5-10 random fields per sample.

#### RNA extraction and Quantitative Real-Time PCR

RNA was prepared from frozen liver samples (~50 mg) by homogenization in Trizol Reagent (Ambion) using a high-speed homogenizer (TissueLyser, Qiagen). Complementary DNA (cDNA) was synthesized from total RNA using the Quanta Biosciences qScript cDNA Synthesis Kit for mRNA, or the Quanta Biosciences qScript microRNA cDNA Synthesis Kit for miRs. Gene expression levels were quantified by qPCR using a Quanta Biosciences SYBR Green PCR Kit with the following primer sets and normalized to Hprt for mRNAs and Rnu6 for miRs.

| Gene | Sense | Anti-sense |
|---|---|---|
| *Apoe* | ttgctgacaggatgcctagc | gtaatcccagaagcggttcag |
| *Cd11β* | gggaggacaaaaactgcctca | acaactaggatcttcgcagcat |
| *Cd68* | tgtctgatcttgctaggaccg | gagagtaacggcctttttgtga |
| *Cpt-1* | agacaagaaccccaacatcc | caaaggtgtcaaatgggaagg |
| *Cyp4a14* | tcagtctatttctggtgctgttc | gagctccttgtccttcagatggt |
| *Cyp4a10* | tccagcagttcccatcacct | ttgcttccccagaaccatct |
| *F4/80* | ccccagtgtccttacagagtg | gtgcccagagtggatgtct |
| *Fasn* | cccctctgttaattggctcc | ttgtggaagtgcaggttagg |
| *Gck* | aggcacgaagacatagacaag | ggagaagtcccacgatgttg |

| | | |
|---|---|---|
| *G6pc* | ggtcacttctactcttgctatctttc | cccagaatcccaaccacaag |
| *Hprt* | gcgatgatgaaccaggttatga | atctcgagcaagtctttcagtcct |
| *IL-6* | agttgccttcttgggactg | cagaattgccattgcacaa |
| *IL-10* | ggttgccaagccttatcgga | acctgctccactgccttgct |
| *miR-21* | tagcttatcagactgatgttga | perfecta® universal pcr primer (cat. no. 95109) |
| *miR-122* | tggagtgtgacaatggtgtttg | perfecta® universal pcr primer (cat. no. 95109) |
| *Pepck* | gacattgcctggatgaagtttg | tggcatttggatttgtcttcac |
| *Pparα* | aacctgaggaagccgttctgtgacat | gaccagctgccgaaggtccaccat |
| *TLR4* | ttcagaacttcagtggctgg | tgttagtccagagaaacttcctg |
| *TNFα* | gaaaagcaagcagccaacca | cggatcatgctttctgtgctc |
| Rnu6 | cgcaaggatgacacgcaaattc | perfecta® universal pcr primer (cat. no. 95109) |

#### Flow Cytometry

FACS analysis was performed on cells freshly isolated from white adipose tissue (WAT) or liver tissue samples from mice anesthetized with ketamine/xylasine following perfusion with PBS essentially as described (Cho et al., 2014) and digested with Liberase® (Roche). For isolation of the stromal vascular fraction from WAT, Liberase digestion of adipose tissue was followed by incubation for 10 minutes with 10mM EDTA, passage through a 100µm filter, and centrifugation. For hepatic tissue, Liberase digestion was followed by 70µm filtration, and centrifugation through a gradient of percoll (GE Healthcare Bio-Sciences) to separate immune cells from parenchymal cells. Following centrifugation, cells were suspended in a red blood cell lysis solution (0.155 M NH4Cl, 0.01 M KHCO3, 0.01 mM EDTA; pH 7.4) for 1 minute, followed by centrifugation and reconstitution in PBS. Fluorochrome-conjugated antibodies against the following antigens were used: anti-mouse-F4/80-PE (eBioscience), anti-mouse integrin αM-FITC (R&D), and anti-mouse cd45-PC5.5A (eBiosience).

#### Taxonomic Microbiota Analysis

Frozen fecal samples were processed for DNA isolation using the MoBio PowerSoil kit (Quiagen) according to the manufacturer's instructions. For analysis of fecal microbiome from mice colonies maintained in the Jerusalem, the 16S rRNA gene PCR amplification, 1 ng of the purified fecal

DNA was used for PCR amplification. Amplicons spanning the variable region V3/4 of the 16S rRNA gene were generated by using the following primers: Fwd 5'-GTGCCAGCMGCCGCGGTAA-3', Rev 5'- GGACTACHVGGGTWTCTAAT-3'. The reactions were subsequently pooled and cleaned (PCR clean kit, Promega), and the PCR products were then sequenced on an Illumina MiSeq with 500 bp paired-end reads. The reads were then processed using the QIIME analysis pipeline. In brief, fasta quality files and a mapping file indicating the barcode sequence corresponding to each sample were used as inputs, reads were split by samples according to the barcode, taxonomical classification was performed using the RDP-classifier, and an OTU table was created. Closed- reference OTU mapping was employed using the Greengenes database. Rarefaction was used to exclude samples with insufficient count of reads per sample. Sequences sharing 97% nucleotide sequence identity in the 16S region were binned into operational taxonomic units (97% ID OTUs).

For analysis of fecal microbiome from mice colonies maintained in Kiel and Hamburg, extracted genomic DNA was used to amplify the 16S rRNA gene specific variable region $V_{3–4}$. Success of 16 S rRNA gene amplification performance was confirmed by running an aliquot of PCR product on a 2% agarose gel. The amplicon quantities were normalized using the SequalPrep™ Normalization Plate Kit (Invitrogen), amplicons were pooled to make a library and sequenced using the MiSeq Reagent Kit v3 (Illumina) at Institute of Clinical Molecular Biology, Kiel, Germany.

Analysis of sequence reads was performed using an in-house shell script pipeline based on standard procedures for 16S rRNA gene sequence data (Kozich et al., 2013). In brief, the multiplex identifiers (MID) and 16S rRNA gene specific amplification primer sequences were removed prior to further sequence analysis. Subsequently, quality-control of sequence reads were performed as defined in Miseq SOP pipeline. Furthermore, sequences were aligned against mothur curated SILVA reference database in Mothur (Schloss et al., 2009). Sequence reads not aligned against 16S rRNA gene V3–V4 (Kiel data sets) or V4 regions only (Jerusulam data sets) were removed from subsequent analysis. Chimeric sequences were detected by the chimera Vsearch algorithm and were also removed. In the first step, sequences were classified (threshold 80%) phylogenetically using mothur formatted greengenes (gg_13_8_99) training sets and eliminated if classified as unknown, archaea, eukaryotes, chloroplast, or mitochondria. Subsequently, reference-based (green genes) operational taxonomical units (OTUs) picking approach was implemented to cluster sequences with same phylogenetic affiliations into a phylotype at genus and phyla level (label = 1). Alpha diversity indices were calculated by Mothur. Further β-diversity estimates and Non-parametric permutational multivariate analysis of variance (NPMANOVA) was performed in PAST (Hammer et al., 2001) for ascertaining the significance of clustering in sampling groups for β-diversity.
